# Macroevolutionary Stability Predicts Interaction Patterns of Species in Seed Dispersal Networks

**DOI:** 10.1101/2020.04.15.042754

**Authors:** Gustavo Burin, Paulo R. Guimaraes, Tiago B. Quental

## Abstract

Assessing deep-time mechanisms affecting the assembly of ecological networks is key to understanding biodiversity changes on broader time scales. We combined analyses of diversification rates with interaction network descriptors from 468 bird species belonging to 29 seed-dispersal networks to show that bird species that contribute most to the network structure of plant-frugivorous interactions belong to lineages that show higher macroevolutionary stability. This association is stronger in warmer, wetter, less seasonal environments. We infer that the macroevolutionary sorting mechanism acts through the regional pool of species by sorting species based on the available relative differences in diversification rates, rather than absolute rates. Our results illustrate how the interplay between interaction patterns and diversification dynamics may shape the organization and long-term dynamics of ecological networks.

## Main Text

Seed dispersal by animals is a fundamental property of ecosystems (1). This mutualism between angiosperms and, mainly, vertebrates started about 80 Mya (2), and currently between 70-90% of woody species rely on vertebrates to disperse their seeds (3). Accordingly, many vertebrate groups have fruits as part of their diet (56% of bird families, and many species of mammals, amphibians, reptiles and fishes – 3–4). The radiations of seed dispersing birds and mammals are hypothesized to be linked to the rise and dominance of angiosperms during the Cenozoic (2), although the causal links are elusive. In particular, some bird groups are consistently recognized as specialized frugivores across broad range of spatial and temporal scales (1, 5–7).

Most studies on seed-dispersal networks have focused on understanding patterns and processes at ecological timescales (e.g. 8–9), with few studies looking at broader temporal scales (2, 10–14). We are now beginning to understand how diversification dynamics may affect the assembly process, and consequently, the structure of interaction networks (e.g. 15–16). Although interaction networks as a whole might be plastic both in time and space (17–19), evidence suggests that the core of seed dispersal networks is robust to yearly fluctuations of fruit availability and bird species presence (20). Moreover, theory suggests that species that show higher persistence across time and space should interact with more partners (21–22), supporting the idea that at least the core of the network might show some temporal stability. This temporal stability in interaction patterns may be observed at longer temporal scales, since closely related frugivores often interact with similar coteries of plants (12, 23). In this context, a fundamental problem to solve is if and how patterns of interaction observed in the networks formed by plants and their seed dispersers at ecological scales are associated with macroevolutionary dynamics operating at deeper temporal scales.

Here, we explore this problem by integrating macroevolutionary information, ecological data on species interactions, and network analysis. We hypothesize that lineages that contain long-lived species or lineages that can quickly accumulate many species of birds (hereafter macroevolutionary stable lineages) are more likely to contribute with species to the core of ecological networks, providing an explicit macroevolutionary mechanism for network assemblage (see Fig. 1 for schematic version of general approach and data used).

**Figure 1:**
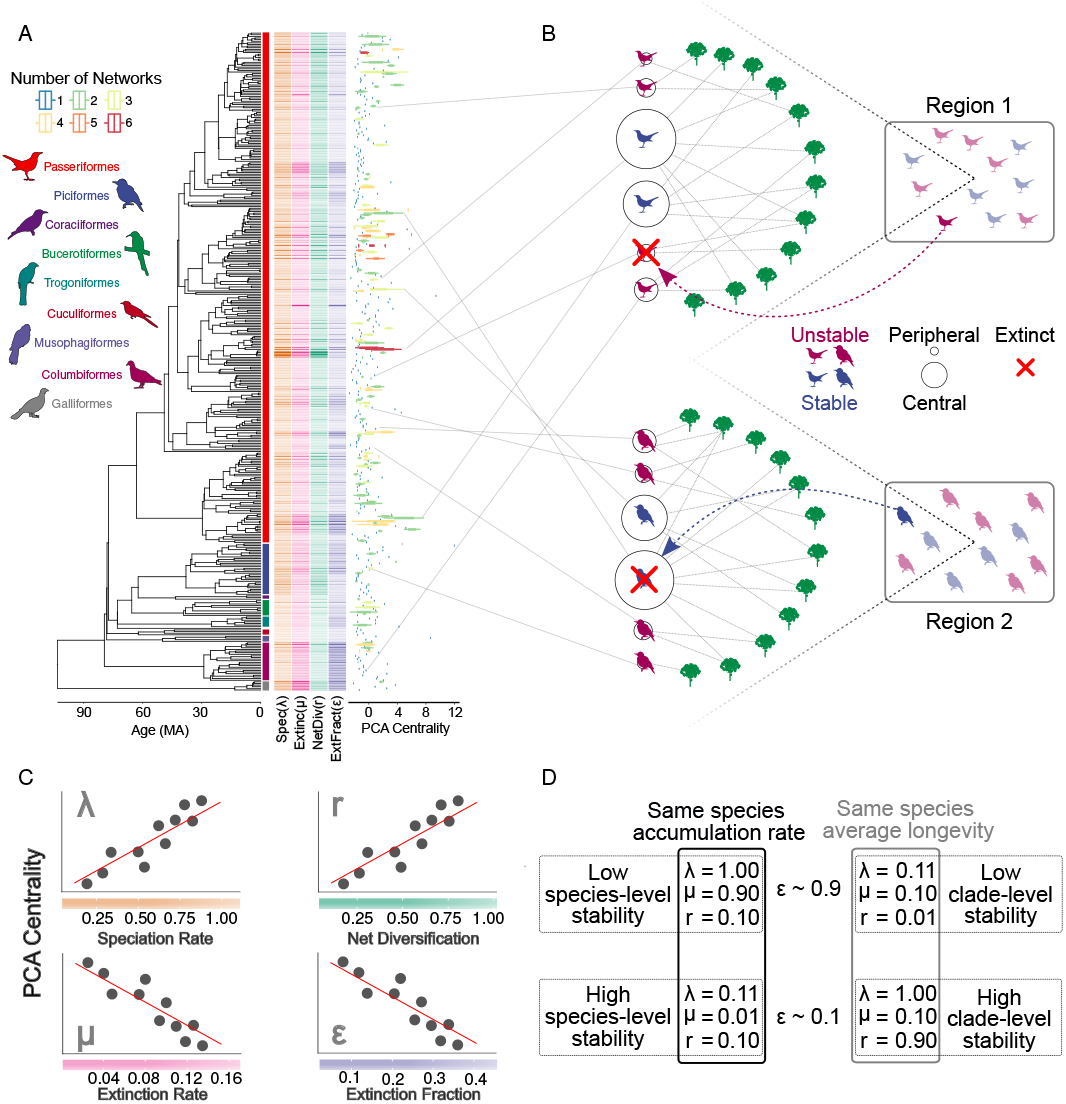
Conceptual framework and data used. A Exemplar pruned phylogeny with indicated bird orders, along with the speciation rates (λ - orange dashes), extinction rates (*μ* - pink dashes), net diversification rates (r - green dashes), extinction fraction (*ϵ* - purple dashes), and the PCA centrality values for each of the species of all species used in the analyses. PCA centrality scores resulting from defining interaction patterns when using three different network metrics. Some species were present in more than one ecological network, and this is information is displayed by points or bars of different colors in the PCA centrality graph. The color scheme on the X-axis on panel C acts as legend for the rate values on panel A.B. Cartoon example of two networks, showing the suggested patterns of network assembly with respect to macroevolutionary rates shown in Panel A, and the potential replacement from another species in the regional pool of species (indicated by the dashed line) in case a given species in a particular network goes extinct. C. Expected correlations between PCA centrality and all four rates considered in this study. D. illustrative combination of speciation and extinction rates showing different degrees of species- and clade-level stability. Accumulation rate is described by the rate of diversification, while average species longevity is the reciprocal of extinction rate (1/*μ*).

Here we define macroevolutionary stability at two different levels. At the species level, we define stable species as species that typically last longer (lower extinction rates imply longer longevities) or species that are more likely to produce new daughter species (those species with higher speciation rates). In the latter case, the continuation of the “species” amid extinctions would be through the production of daughter species that likely share traits associated with the interaction (note that this case could be also seen as a higher-level effect – see below). At higher phylogenetic levels (e.g., monophyletic lineage with multiple species), we define stable lineages as lineages that either have low extinction fraction (i.e., relatively low extinction compared to speciation), and/or higher net diversification (i.e., lineages that can accumulate species at a faster pace), which would allow more efficient replacement of a given extinct species by a closely related one that would show similar patterns of interaction (12, 23).

Starting from a molecular phylogeny (Fig. 1A), we estimated rates of speciation, extinction, net diversification rate (speciation minus extinction), and extinction fraction (extinction divided by speciation rate) for all species in the networks. We then estimated species’ interaction patterns (how they were connected) within each of the 29 networks (Fig. 1B) using three different network descriptors to characterize interaction patterns of each species which were then combined into a single descriptor index for each species by using principal component analysis - PCA (24). Lastly, we used a hierarchical Bayesian phylogenetic framework to test for relationships between macroevolutionary stability and interaction patterns of bird species (Fig. 1C) (24). This workflow naturally incorporated phylogenetic and ecological uncertainties, along with environmental factors, to address how the interplay between biotic and abiotic factors affect the assembly of local networks according to macroevolutionary dynamics. We jointly tested whether the sorting of different macroevolutionary diversification dynamics (speciation and extinction rates and combinations of these two rates, see Fig. 1) takes place at the regional or global scale by using raw absolute rates and standardized rates (re-scaled for each network). The absolute rates allow us to explore a global sorting mechanism which selects for specific diversification dynamics (specific absolute rate values) irrespective of the pool of available clades. Conversely, standardized rates allow us to explore a sorting mechanism that would act on the relative rank of rates available at the regional scale. Because of the lack of comprehensive species-level phylogenies for plants, we performed analysis only for bird species.

We found that central bird species in seed dispersal networks tend to belong to macroevolutionary stable lineages. Standardized speciation and extinction rates show respectively positive and negative correlations with species’ patterns of interaction (Fig. 2A). The negative correlation between interaction pattern and extinction rates highlights that the sorting process takes place at the species level (because longevity is an inherent property of a species), with longer-lasting species occupying more central roles positions within networks. We also found that extinction fraction is negatively correlated with species’ interaction patterns while net diversification rates are positively correlated (Fig. 2B). In all cases the posterior distributions are not centered on a slope value of zero. These correlations suggest that species that occupy central positions in networks tend to belong to lineages that are either less volatile (sensu 25) and/or that generate multiple species in a short time span. Hence central species are both more likely to persist in time (negative correlation with extinction rate), and to belong to clades that are more likely to provide a replacement species if one goes extinct (correlations with extinction fraction and diversification rate).

**Figure 2:**
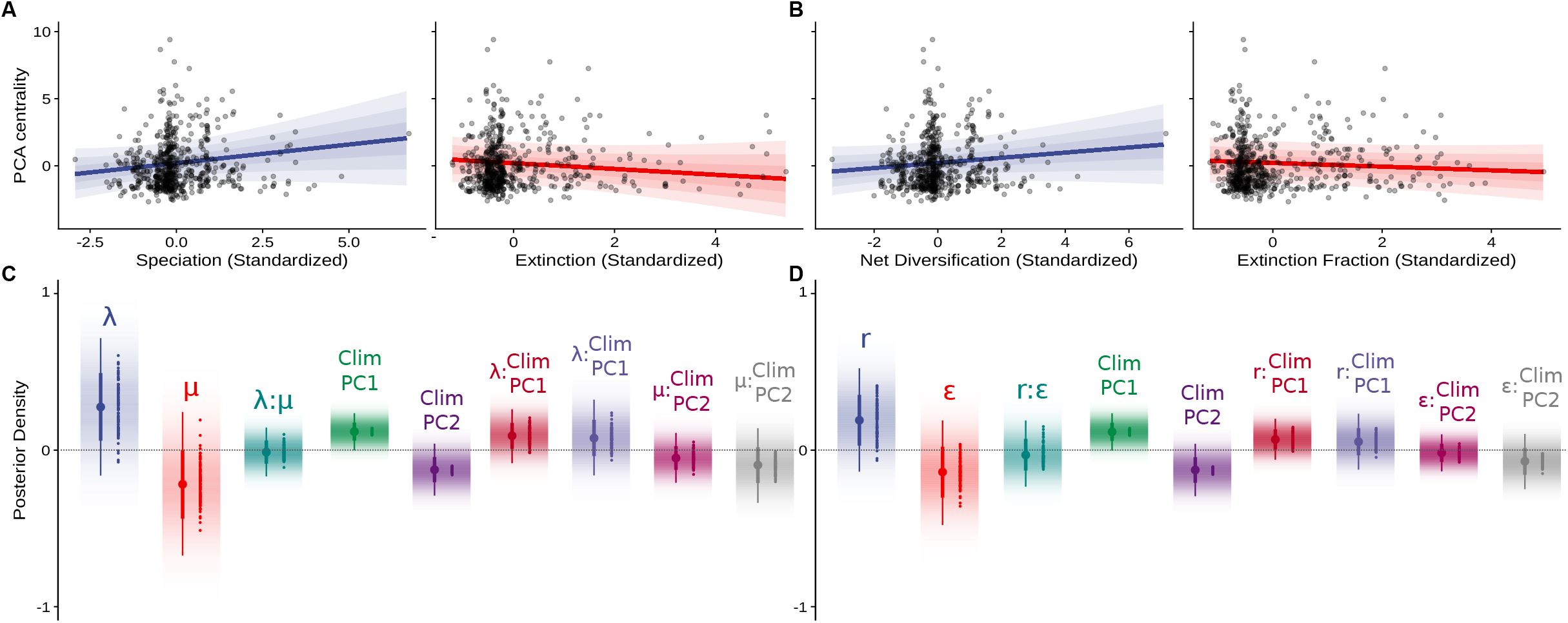
Association between network centrality and macroevolutionary stability. A. Exemplar association between PCA centrality values (a PCA score based on three centrality metrics) and speciation and extinction rates estimated from one phylogenetic tree. The blue and red lines are the median intercept and slope values from the combined posterior distribution of each parameter. B. Exemplar association between PCA centrality scores and extinction fraction and net diversification rates estimated from one phylogenetic tree. The blue and red lines are the median intercept and slope values from the combined posterior distribution of each parameter. Note that in A and B the points are non-independent due to phylogenetic structure and therefore visual inspection of results could be misleading. C. Posterior distributions of all slopesfor Bayesian generalized linear mixed model analysis between using speciation (λ) and extinction (*μ*) rates as predictors. D. Posterior distributions of all slopes) for the Bayesian generalized linear mixed model analysis between using net diversification (r) and extinction fraction (*ϵ*) rates as predictors. Posterior distribution of most slopes are off-centered from a slope of zero (positive for speciation and net diversification, and negative for extinction and extinction fraction), indicating that species belonging to macroevolutionary more stable lineages are more central in the networks. In panels C and D the density of color represents the posterior density of each parameter; the point-and-range lines on the left of each variable represents the median (point), 66% highest posterior density interval (HPD - thick line) and 95% HPD (thin line); the dots on the right of each variable represent the median of the posterior distribution of each individual tree used in the analyses. ClimPC1 and ClimPC2 refer to the principal component of climatic variables, with ClimPC1 mainly representing variation in average annual temperature and temperature seasonality, and ClimPC2 representing total annual precipitation and precipitation seasonality (see Fig S9 for the loadings).

We also found a positive relationship between speciation rate and centrality measures, reinforcing the idea of a clade-level stability mechanism because high speciation rates indicate that those species are more likely to produce daughter species that might replace it if it goes extinct. One important assumption for the higher-level sorting mechanism, which finds support in previous knowledge (12), is that the replacement species has similar ecological attributes that allow them to perform similar ecological dispersal services. All these results hold after accounting for multiple sources of uncertainty (Fig. S1 and S2). Median standardized effects for speciation, extinction, extinction fraction and net diversification were 0.275, −0.218, −0.140 and 0.191, respectively, indicating the two higher-level components of macroevolutionary stability contribute with similar intensity to the structuring of the networks. Sensitivity analysis based on the medians of the posterior distributions for most parameters for each individual tree (Fig. 2C, 2D) suggests that this signal holds irrespective of phylogenetic uncertainty.

Our results provide evidence that this macroevolutionary sorting of diversification dynamics is predominantly a regional-scale phenomenon, because we only observed evidence for sorting when analyzing the rates standardized within networks (Fig. 2), but not for the raw absolute rates (Fig. S5 and S6). This suggests that the macroevolutionary sorting mechanism acts at a regional scale sorting the species within each region through their relative rank of stability, rather than on absolute values of speciation, extinction, extinction fraction or net diversification rates. Our results show that representatives of important seed-dispersers groups across multiple localities indeed show high relative macroevolutionary stability either due to fast species accumulation (e.g., thraupid genera such as Tangara and Thraupis), or to long-lived lineages (e.g., species of Turdidae) (5–7). It is also worth noting that ecological factors such as species abundance distributions (26) and the presence of invasive species (27) also influence network organization. Unfortunately, the lack of data prevents us from further testing if macroevolutionary consequences to network structure is modulated by those factors.

To evaluate if species within the same lineage have similar interaction patterns in different networks, we calculated the average centrality value for each different lineage (either family or genus) for all networks. The association between the mean centrality value of each lineage in different networks decays as a function of the geographical distances between those networks (Fig. 3 and Fig. S7), suggesting that geographically close networks tend to have similar lineages with similar interaction patterns, but similar patterns of interaction are shown by different lineages in different places. This reinforces previous findings that different networks along a geographical range are structured with similar functional roles within their structures (28). This result also reinforces that the assembly process occurs at the regional scale, in accordance with existence of a relationship between centrality and macroevolutionary stability for the standardized rates but not for the absolute rates.

**Figure 3:**
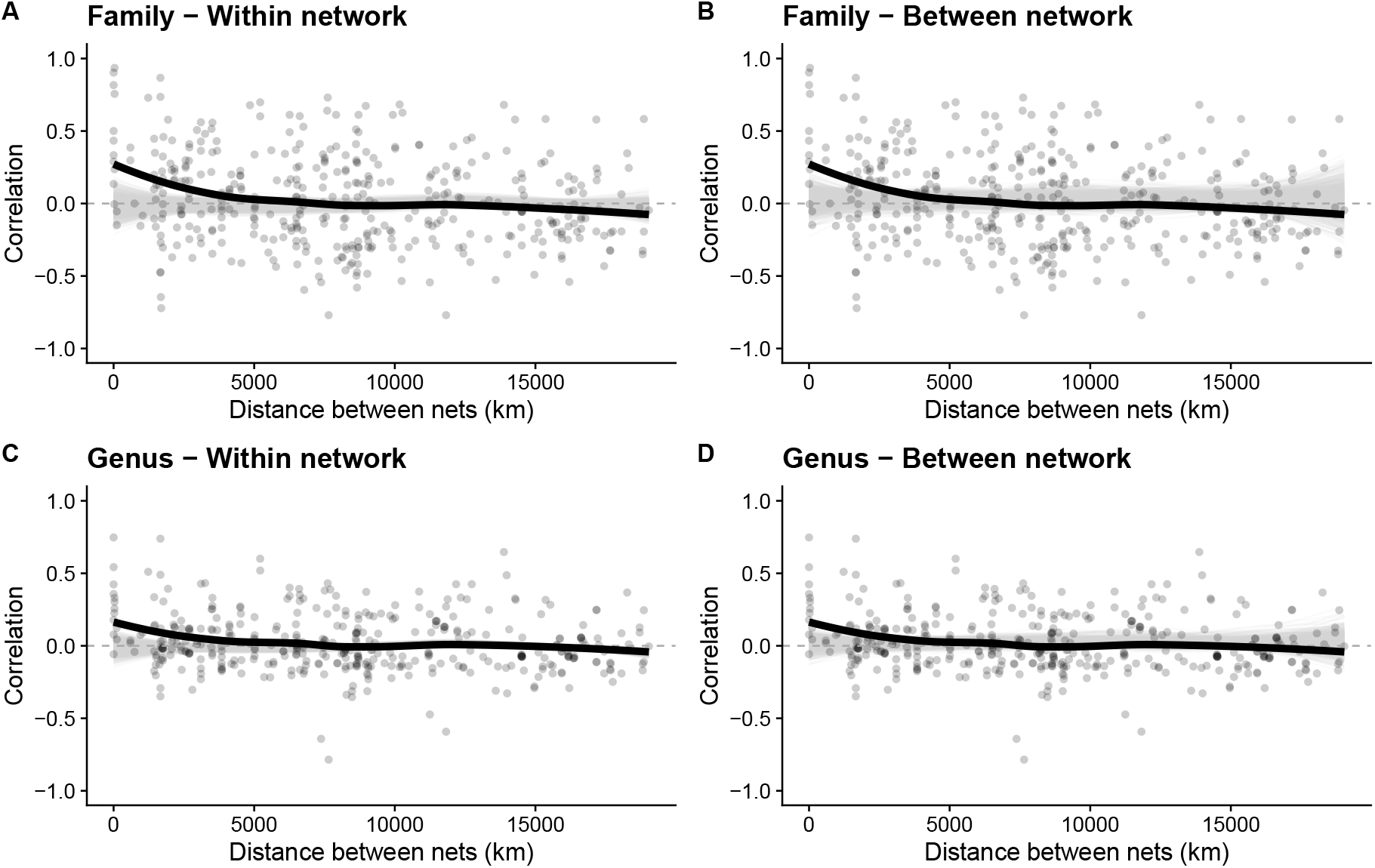
Association between mean PCA centrality for each family (A and B) or genus (C and D) and geographical distance. Each point represents a pairwise correlation between interaction patterns in two different networks, and the red lines are a loess smoothing showing the general trend of the data. The grey areas represent the null models that were built by randomizing centrality values for each species within networks (panels A and C), or by randomizing the identity of networks for each geographical distance (panels B and D). Regardless of the null model selected, the associations between mean centrality of families/genera are higher than expected by the null model for geographically close networks, and this similarity dissipates with distance, suggesting there is no single group driving the association between macroevolutionary stability and interaction patterns.

We also found that environmental conditions affect the relationship between interaction patterns in the seed dispersal network and macroevolutionary stability, with warmer, wetter, non-seasonal environments showing a stronger sorting (here revealed by an interaction with environmental descriptors, which suggests a steeper slope between macroevolutionary rates and centrality descriptors), favoring the building of the network around macroevolutionary stable species and lineages (Fig. 2). These environments harbor the highest frugivorous species richness (7), and we hypothesize that such species-rich environments allow for a finer sub-division of network roles on which this macroevolutionary sorting of stable species and lineages can act. In fact, networks in these tropical-forest-like environments often show higher variation in centrality values across species than networks found on colder, drier, and seasonal environments (Fig. 2C, 2D). This variation, however, was not simply a consequence of variation in species richness. Body size disparity data show that niche space in warmer, more humid environments is more homogeneously occupied without increasing neither total niche space nor niche overlap (using body size as a proxy) between species (Fig. S10 and S11). Furthermore, we found that not only central species interact with more partners, but also these partners belong to more distinct phylogenetic groups (Fig. S12-S15). Thus, higher variation in species richness, niche occupation and diversity of partners provide the raw material that allows macroevolutionary sorting of stable species to occur.

Our results provide evidence for a macroevolutionary sorting mechanism (species selection in a broad sense - 29) on network assembly where central species tend to have longer longevities (the inverse of extinction rate) and belong to evolutionary lineages that are more stable over deep time. Moreover, we found that central species not only interact with a higher, more (directly or indirectly) connected number of species, and that those partner species belong to more distinct ecologies (Fig. S12-S15). Although the relationship between macroevolutionary rates and network centrality metrics may reflect the role of macroevolutionary stability on the sorting of interaction patterns during network assembly, we note that the causal relationship might also act in the opposite direction, and species network position might eventually affect the rates of diversification. For instance, increased frugivory may fuel diversification within some vertebrate lineages, such as primates (30). We are only beginning to understand how macroevolutionary effects relate to the ecological organization of species interactions networks. In this sense, it is hard to infer causality direction when experiments are not an option. Nevertheless, our study shows evidence of the importance of species turnover on the structure of ecological networks in geological time, expanding the temporal scale to the ones addressed in previous studies (17, 19). By now, our results suggest potential multi-level selective regimes involving interaction patterns and diversification dynamics, which might shape the fate of groups of very distantly related lineages (e.g., birds and plants) linked through ecological interactions.

## Acknowledgments

We thank Eduardo Santos, Marco Melo, Marcus Aguiar, Patrícia Morelatto, Marilia Gaiarsa, Daniel Caetano, Natalie Cooper, Sarah Alewijnse, Travis Park, and all the researchers in LabMeMe and Guimarães Lab for their insightful comments during several steps of this work.

## Author contributions

Gustavo Burin: Conceptualization, Methodology, Data Curation, Investigation, Writing - Original Draft & Review & Editing, Visualization;

Paulo R. Guimarães Jr.: Conceptualization, Methodology, Writing - Review & Editing;

Tiago B. Quental: Conceptualization, Methodology, Writing - Original Draft & Review & Editing, Supervision;

## Data and materials availability

Data and materials availability: Bird phylogenies were obtained from birdtree.org (33), and ecological networks were compiled by 38 and can be directly downloaded from the following Dryad link: http://dx.doi.org/10.5061/dryad.2br2b. All data and code used in this study is also available for download at https://doi.org/10.5281/zenodo.3560680. Codes are also available at https://www.github.com/gburin/macroevoNet.

