## Supplementary Material for "Macroevolutionary Stability Predicts Interaction Patterns of Species in Seed Dispersal Networks"

### Supplementary Material - Macroevolutionary imprints on the assembly of seed dispersal networks

#### Materials and Methods

To investigate the potential effect of diversification rates on the assembly of current ecological networks we first characterized each frugivore species with respect to its interaction patterns in the network, and then estimated speciation and extinction rates related to each bird lineage using a comprehensive molecular phylogeny containing approximately 67% of all bird species (see below). We then tested for an association between the network descriptor of the interaction patterns and the macroevolutionary rate (see figure 1 main text for a cartoon version of the general approach) using phylogenetic generalized linear mixed models (MCMCglmm - 31). Phylogenetic correction was used because the network descriptors showed significant phylogenetic signal (mean value for Pagel's lambda = 0.537 but note that for each tree the corresponding lambda value was used).

##### Diversification Rates

To estimate speciation and extinction rates we used the most recent bird phylogeny, which comprises the 99.3% of the extant species (9993 species in the phylogeny from an estimated total of 10,064 species – (32). To avoid any potential bias introduced by adding species without DNA sequences (using a pure-birth algorithm) on our estimates of speciation and extinction rates (33), we used the trees that contain only species for which molecular data were available from the original phylogenetic paper (32). This tree had 6670 species, but species not sampled were explicitly taken into account by the model. We analyzed the bird molecular phylogeny using a Bayesian framework (BAMM v. 2.5; 34) to estimate the rates of speciation and extinction while explicitly taking into account the sampling fraction of species. The incompleteness (about 1/3 of species without DNA data) was considered within BAMM by using the corresponding percentage of missing species within each clade of the maximum clade credibility tree obtained by Jetz et al. (32). Rabosky (34) has developed an algorithm that finds sub-trees within a phylogeny, which share speciation and extinction rates through a Bayesian framework using reversible-jump Markov Chain Monte Carlo sampling. Each diversification shift configuration has an associated posterior probability, and each configuration can assign different diversification rates to a particular species or clade. Therefore, the rates were estimated using the whole tree, and we then obtained the mean rates averaged by the marginal posterior probabilities of each distinct shift configuration for each species to be used in the following steps, thus accounting for the uncertainty related to the true rate regime configuration. This way, we also avoid any issue related to rate estimation for independently extracted subclades from a tree (35).

In addition to the traditional rates obtained from BAMM ($\lambda$ and $\mu$, speciation and extinction rates respectively), we calculated other two of our rates of interest: the extinction fraction ($\varepsilon = \mu/\lambda$) and net diversification rate ($r = \lambda - \mu$) for all species in the networks. These two rates represent two different metrics used to quantify the stability/predictability of a given lineage, instead of characterizing the macroevolutionary dynamics of each single species like the probability of speciation and extinction. A higher value of extinction fraction suggests a more volatile dynamics within a given lineage and hence a higher chance of the entire lineage going extinct. A lower net diversification rate suggests that a lineage accumulates species as a smaller pace, and therefore has a higher chance of going extinct than lineages with higher net diversification rate. To our purposes, we build two sets of models that aim to assess the relationship between ecological interaction pattern and macroevolutionary stability at different levels: one that uses speciation and extinction rates (for a species-level sorting process) and one that uses extinction fraction and net diversification (for a clade-level sorting process). We assume that the two rates of each set of models (each of them plus their interaction) affects the likelihood that a given lineage might be evolutionary stable/reliable at long temporal scales. In all analyses, we used rates estimated at the present, as it coincides with the timeframe of the data for the networks, and also because these rates were shown to be both well estimated with BAMM (36), and not so affected by parameter identifiability issues (37).

To evaluate the geographical scale in which the sorting process takes place, we replicated the two sets of tests using both the raw rates and rates standardized by network. To standardize the rates we used a z-score procedure: we re-scaled the rates by calculating the difference between the rate of each species and the mean rate of the network, and then dividing this difference by the standard deviation of the rates in the network. With this z-score transformation, we test if the sorting acts based on the global (absolute) rate of a species or if it acts on the rate relative to other species available in the regional species scale.

#### Ecological Networks

To evaluate the ecological interaction pattern of different species we used data from 34 different frugivory networks compiled by previous authors (38). Those networks comprise both temperate and tropical areas and a total of 546 unique bird species (838 total unique interactions). From the original dataset, we discarded 5 networks due to biases in species composition (networks that focused on specific bird or plant groups, and therefore representing a small subset of the interacting assemblage), resulting in a final dataset of 468 species (700 unique interactions).

Although some networks (23) had information on the strength of interactions, we decided to binarize all networks for two reasons. First, it allows us to work with a larger dataset. Second, it is known that interaction strengths vary across time (17-19), rendering quantitative estimates not necessarily meaningful as a long-term description. We also removed pairs of species that only interacted with each other, and were, by definition, disconnected from the rest of the network. Those represent a small set of species since all networks show a giant component (39), i.e., the vast majority of species are connected to each other through direct or indirect pathways. Indeed, only 14 species from only 6 different networks from a total of 468 species from 29 networks. This was done because some of the network descriptors are not defined for networks that are not connected.

To characterize each species' interaction patterns within the compiled dataset we investigated different node (species) properties of the networks by calculating three different centrality descriptors using the original bipartite networks: degree (number of interactions), closeness centrality (the reciprocal of the sum of shortest distances, in links, connecting the focal node to all other nodes in the network) and Katz centrality (a measure of the distance, in terms of all possible pathways, between the focal node and all other nodes of the network, using $\alpha = 0.05$). Such metrics provide us with information about the importance of each species network structure when considering only the direct interactions (degree), the shortest pathways (closeness) or all pathways (Katz). To be able to combine the interaction patterns estimated for different networks (that have different properties, such as number of species and connectance) into a single analysis, we first standardized the three metrics by calculating their z-scores for each network separately. After compiling each of the three standardized centrality metrics for all species in all networks in a single dataset, we performed a Principal Component Analysis (PCA) in order to obtain a single score that comprised different aspects of the network position (as a proxy for the interaction patterns - i.e. ecological role). We used only the first Principal Component since it explained 93.3% of the variance in centrality values.

#### Sources of uncertainties

We incorporated in our analyses two additional distinct sources of uncertainty (besides BAMM's rate configuration uncertainty previously mentioned). The first one is related to the true evolutionary history of birds. To incorporate the phylogenetic uncertainty, we used 112 different topologies from the bird phylogeny (56 of each backbone) to both estimate the rates of diversification and to take into account the phylogenetic structure of the residuals in the linear model. This number of trees is more than double the average recommended number of trees to encompass enough phylogenetic uncertainty in comparative analyses (40). The other source of uncertainty came from the networks. From all the species in the final dataset (N = 468 species), 139 are part of more than one network (up to 6 networks), making it impossible to attribute a single centrality value to those species. We thus incorporated this variation as a source of uncertainty in the regression analysis by using species identity as a random factor as described in the modeling section below.

#### Co-factors

Because the environment at which species are found might modulate the nature and relevance of ecological interactions (41), we ran a global analysis (with all networks combined) using climatic variables as co-factors. Those were: 1 - annual mean temperature; 2 - temperature seasonality; 3 - annual precipitation; 4 - precipitation seasonality. To summarize environmental conditions and avoid over-parameterization of the models, we ran a PCA with the four climatic variables, and used the first and second PCs that in combination explain 81.92% of total variance. We note that those two PCs aligned well with the raw temperature and precipitation variables (Annual Mean Temperature, Temperature Seasonality, and Annual Precipitation have scores of 0.619, -0.592, and 0.492 on PC1, respectively. Annual Precipitation and Precipitation Seasonality have scores of 0.331 and -0.942 on PC2, respectively - Fig. S8). The climatic data were obtained from the WorldClim database version 2.0 (42).

#### Modelling

We tested the association between interaction patterns and macroevolutionary dynamics using a Bayesian generalized linear mixed model (MCMCglmm), implemented in the R package MCMCglmm (31). This framework allows us to naturally incorporate both sources of uncertainty. By using species identity and the phylogenetic structure as random factors, we can account for intraspecific variation in the interaction patterns on the estimation of the regression parameters while simultaneously controlling for the phylogenetic structure. Moreover, by combining the posterior distributions of parameter values using rates from different trees (topologies), we also can account for phylogenetic uncertainty. We assessed the effect size of each variable used in the model by analyzing the asymmetries in both the combined posterior distributions in relation to zero (no association), and in the medians of individual posterior distributions for each tree (43, 44). By doing this we embrace sources of uncertainty incorporated in our analysis (ecological and phylogenetic) while looking at the magnitude of the effect (43). All MCMC chains were run for 5000000 iterations, with half of the total length discarded as burn-in, and sampling every 2500 iterations. We used inversed Wishart distributions as priors for both the fixed and the random effects. The models we tested were the following:

Speciation and Extinction:

PCA Centrality ~ $\lambda$ + $\mu$ + $\lambda : \mu$ + ClimPC1 + ClimPC2 + $\lambda$:ClimPC1 + $\lambda$:ClimPC2 + $\mu$:ClimPC1 + $\mu$:ClimPC2

Net Diversification and Extinction Fraction:

PCA Centrality ~ r + $\epsilon$ + r:$\epsilon$ + ClimPC1 + ClimPC2 + r:ClimPC1 + r:ClimPC2 + $\epsilon$:ClimPC1 + $\epsilon$:ClimPC2

We fitted the two models using both the raw rates and the standardized (z-scored) rates. Additionally, we also fitted the same two models for the standardized rates but excluding species in the top 5% standardized rates to check for possible biases on the final results (Figs. S4 and S5).

#### Geographical patterns of centrality per family/genus

To check if the species that play central/peripheral roles in different networks show a spatial structure with respect to their taxonomic identity (e.g. belong to the same family/genus), we calculated the association between the average centrality per family or genus for each pair of networks using Spearman's correlations, ρ, and then plotted ρ as a function of the distance between each pair of networks. We used family and genus as our data points (two separate analysis) for it is unlikely that the same species would be present in geographically distant networks. To build a null model, we used two different randomization approaches and two different ways to deal with missing data. Hence, we ended up with four different null models for each analysis, one pair for the family and another for the genus level.

In the first randomization approach (Fig. 3A and C), we randomized the species centrality values within each network prior to calculating the average centrality for each lineage (either per family or per genus, two separate analysis each using one taxonomic rank). The association was then estimated for all pair of networks and plotted as a function of the geographical distance of each network pair. In the second randomization approach (Fig. 3B and D), instead of randomizing the species centrality values within each network, we randomized the network's geographical location by randomly sampling the latitude/longitude for each of the networks from the pool of coordinates of the entire dataset without replacement, recalculated the pairwise association of average centrality, and plotted these as a function of geographical distance between each pair of networks. Hence in our first randomization approach the species identity is randomized but the geographical distances between pairs of networks is kept as in the empirical dataset. In the second randomization approach it is the geographical distance that is randomized while keeping the original species identity.

In both randomization approaches, there were instances where a given lineage was only present in one of the two comparing networks. Hence to calculate the association between networks when using lineage (e.g. family or genus) average centralities, we had to deal with the limitation that "valid" average centrality value for a given lineage might not be estimated for some of the networks. To overcome this limitation, we filled the missing values in two different ways. In the first, we simply replaced the missing values in a given network by 0 (hereafter called "single 0 scenario"). By doing this we are strongly "breaking" any possible association, which means that only very strong association would still remain significant after adding those zero values (Fig. S7). The second approach consisted in replacing the average centrality of a lineage (family or genus depending on the analysis) by 0 not only for the network where that lineage was missing but in both networks ("full 0 scenario"). This was done in order to "completely remove" the effect of that lineage from the analysis, while still allowing for the association between the families/genera that are present in both networks to be estimated (Fig. S7). The prediction of the null model is constructed by doing a LOESS smoothing to each randomized dataset (1000 datasets in total). The empirical LOESS smoothing line is then compared to the group of LOESS smoothing lines defining the null model and the relationship is considered significant if the empirical line does not overlap with the null model lines.

#### Ecospace Occupation

To analyze how ecospace is occupied in different climates, we used an independent dataset of body size information (EltonTraits v1.0, 45) to which we did not perform any curatorial work. Here we assume that body size is a good proxy for ecology (46), but we note that all species used here consume fruits in their diet (they are the species found in the networks). We calculated three different disparity metrics for each network using euclidian distances between body mass values of pairs of species: amplitude (that indicates how broad the niche is for the whole network), average pairwise distance and mean nearest-neighbor distance (that indicates, on average, how far each species is from the closest species in their respective network). We tested the relationship between each of the three metrics with Climatic PC1 (which explains about 56% of the variation - Fig S8) to assess any trends of ecospace occupation along climatic gradients. Those comparisons were done using the disparity metric directly or using the residuals between the disparity metric and species richness. The later analysis was done to control for the effect that richness has on the disparity metric (both are correlated).

#### Degree of interaction generalization

For each bird species in each network, we calculated both the number of unique families with which they interact, and also the plant phylogenetic diversity by summing all branch lengths in the pruned family-level phylogeny (47) containing only the plant families with which the species interact. In order to test if more central species are more generalist (either defined as interacting with a higher number of plant families or by a higher value of plant phylogenetic diversity of their plant partners) we fitted linear models to test for the association between interaction patterns (PCA centrality) and both the number of families and phylogenetic diversity. We have also removed possible effects of both the "number of species" in "the number of families", and of "the number of families" in the "phylogenetic diversity" by using the residuals of these two correlations to test for associations with PCA centrality.

#### Network metadata

Table S1 contains all relevant information of the network dataset used in the main paper. These networks were compiled by Pigot et al. 2016.

Table S1: Metadata of all networks used in this study. The numbering for location references match the ones from Pigot et al., 2016.

| Network | n.bird.species | n.plant.species | n.interactions | Location | Latitude | lat_dec | long_dec | ann.mean.temp | temp.seas | ann.prec | prec.seas |
|---|---|---|---|---|---|---|---|---|---|---|---|
| BAIR | 21 | 7 | 655 | Princeton, USA [1] | Temperate | 40.331800 | -74.66810 | 11.5 | 865.9 | 114.2 | 1.4 |
| BEEH | 9 | 31 | 1189 | Mount Missim, New Guinea [2] | Tropical | -7.266667 | 146.70000 | 19.8 | 59.1 | 216.7 | 2.5 |
| CACG | 15 | 25 | 230 | Caguana, Puerto Rico [3] | Tropical | 18.295000 | -66.78111 | 22.7 | 120.7 | 227.6 | 4.2 |
| CACI | 20 | 34 | 478 | Cialitos, Puerto Rico [3] | Tropical | 18.339167 | -66.46889 | 25.0 | 127.4 | 174.8 | 2.4 |
| CACO | 13 | 25 | 122 | Cordillera, Puerto Rico [3] | Tropical | 18.339167 | -66.46889 | 25.0 | 127.4 | 174.8 | 2.4 |
| CAFR | 15 | 21 | 118 | Fronton, Puerto Rico [3] | Tropical | 18.339167 | -66.46889 | 25.0 | 127.4 | 174.8 | 2.4 |
| CROM | 7 | 72 | 143 | Queensland [4] | Tropical | -17.850000 | 146.08333 | 23.7 | 275.2 | 299.8 | 7.7 |
| DOWS | 15 | 120 | 333 | Highlands, Malawi [23] | Tropical | -10.800000 | 33.80000 | 16.8 | 208.4 | 112.8 | 9.8 |
| FROS | 9 | 16 | 3267 | Mtunzini, South Africa [5] | Temperate | -28.959700 | 31.75010 | 21.3 | 270.9 | 108.8 | 3.9 |
| GEN_A | 18 | 7 | 150 | Santa Genebra Reserve T1, Brazil [6] | Tropical | -22.822222 | -47.11111 | 19.8 | 236.9 | 130.2 | 6.7 |
| GEN_B | 29 | 35 | 397 | Santa Genebra Reserve T2, Brazil [6] | Tropical | -22.822222 | -47.11111 | 19.8 | 236.9 | 130.2 | 6.7 |
| GORCH_A | 7 | 37 | 29 | Loreto, Peru [18] | Tropical | -4.916667 | -73.75000 | 26.9 | 42.1 | 260.0 | 2.0 |
| GORCH_B | 6 | 6 | 200 | Loreto, Peru [18] | Tropical | -4.916667 | -73.75000 | 26.9 | 42.1 | 260.0 | 2.0 |
| HAMM | 16 | 29 | 175 | North Negros, Philippines [7] | Tropical | 10.683333 | 123.18333 | 24.3 | 62.4 | 228.7 | 4.0 |
| HRAT | 17 | 16 | 34044 | Hato Rat□n, Spain [8] | Temperate | 37.150000 | -2.65000 | 15.0 | 573.9 | 37.8 | 4.5 |
| KANT | 25 | 5 | 80 | Campeche state, Mexico [9] | Tropical | 18.505400 | -89.39350 | 24.6 | 221.8 | 112.5 | 5.8 |
| KITA | 22 | 12 | 73 | Khao Yai, Thailand [20] | Tropical | 14.166667 | 101.37500 | 25.9 | 173.7 | 149.4 | 7.4 |
| LAMB | 60 | 25 | 504 | Krau Game Reserve, Malaysia [10] | Tropical | 3.716667 | 102.28333 | 25.4 | 49.9 | 221.8 | 2.8 |
| MACK | 29 | 28 | 58 | Crater Mountain, New Guinea [11] | Tropical | -6.723333 | 145.09333 | 21.8 | 60.4 | 348.3 | 1.4 |
| MONT | 40 | 169 | 666 | Monteverde, Costa Rica [12] | Tropical | 10.300000 | -84.80000 | 20.7 | 65.2 | 291.4 | 5.6 |
| NCOR | 33 | 25 | 7010 | Nava Correhuelas, Spain [ P. Jordano, unpubl.] [13] | Temperate | 38.158200 | -2.73820 | 12.1 | 661.3 | 55.7 | 3.8 |
| NNOG | 28 | 18 | 129 | Nava Noguera, Spain [ P. Jordano, unpubl.] [13] | Temperate | 38.158200 | -2.73820 | 12.1 | 661.3 | 55.7 | 3.8 |
| POULI_A | 19 | 4 | 292 | Panama [13] | Tropical | 8.966667 | -79.53333 | 26.7 | 50.1 | 195.1 | 6.2 |
| POULI_B | 11 | 13 | 1415 | Panama [13] | Tropical | 8.966667 | -79.53333 | 26.7 | 50.1 | 195.1 | 6.2 |
| SAAV_A | 36 | 34 | 354 | Andes, Bolivia [21] | Tropical | -16.410306 | -67.52697 | 18.4 | 121.8 | 113.3 | 6.5 |
| SAAV_B | 21 | 20 | 145 | Andes, Bolivia [21] | Tropical | -16.410306 | -67.52697 | 18.4 | 121.8 | 113.3 | 6.5 |
| SAPF | 8 | 15 | 60 | Yakushima Island, Japan [14] | Temperate | 30.344600 | 130.51270 | 16.2 | 562.9 | 335.6 | 4.1 |
| SARM | 8 | 28 | 41 | Atlantic forest, Brazil [22] | Tropical | -8.966667 | -36.05000 | 21.9 | 139.7 | 129.5 | 5.9 |
| SCHLE | 83 | 32 | 154 | Kakamega Forest, Kenya [17] | Tropical | 0.283333 | 34.78333 | 20.6 | 66.3 | 191.7 | 3.7 |
| SOREN | 14 | 11 | 2745 | Wytham Woods, UK [16] | Temperate | 51.776300 | -1.31400 | 9.8 | 466.1 | 63.3 | 1.3 |
| STEIB | 31 | 31 | 844 | Hesse, Germany [13] | Temperate | 50.652100 | 9.16240 | 8.4 | 636.9 | 75.3 | 1.3 |
| VELHO_A | 29 | 6 | 626 | Arunchal Pradesh, India [19] | Tropical | 27.000000 | 92.76667 | 24.1 | 415.3 | 221.8 | 8.6 |
| VELHO_B | 23 | 6 | 700 | Arunchal Pradesh, India [19] | Tropical | 27.016667 | 92.55000 | 20.3 | 435.3 | 215.0 | 8.9 |
| WES | 81 | 178 | 975 | Intervales and Saibadela, Brazil [15] | Tropical | -14.268417 | -48.41381 | 24.1 | 76.5 | 183.0 | 8.2 |

### MCMCglmm

#### Individual posterior distributions per tree - Standardized Epsilon and r

The main figure shows the combined posterior distribution for all 112 trees used in the study. Below the posterior distributions for each tree are plotted individually.

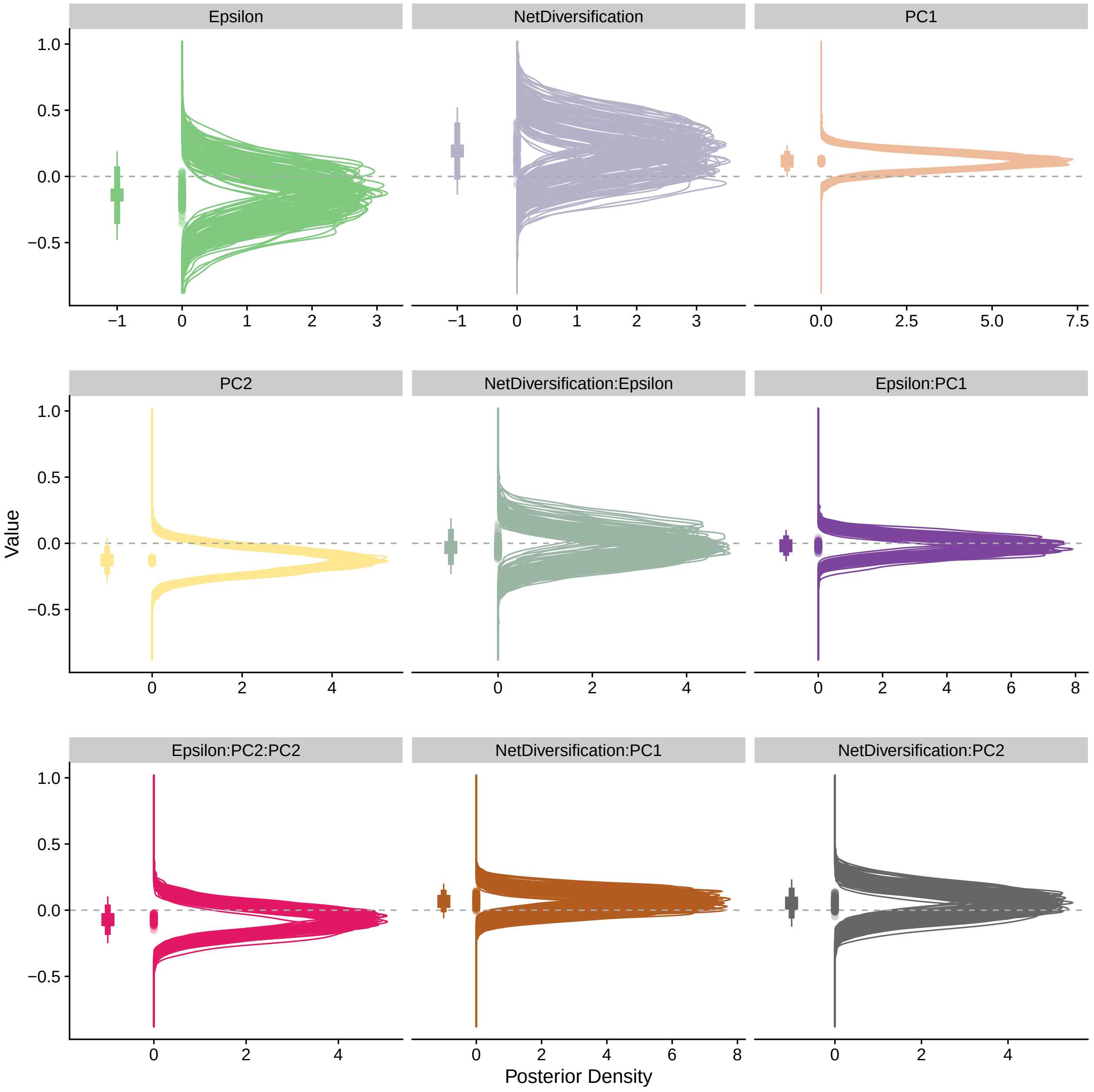


Figure S1: Individual posterior distributions for all trees used using standardized extinction fraction and net diversification rates.

#### Individual posterior distributions per tree - Standardized lambda and mu

The main figure shows the combined posterior distribution for all 112 trees used in the study. Below the posterior distributions for each tree are plotted individually.

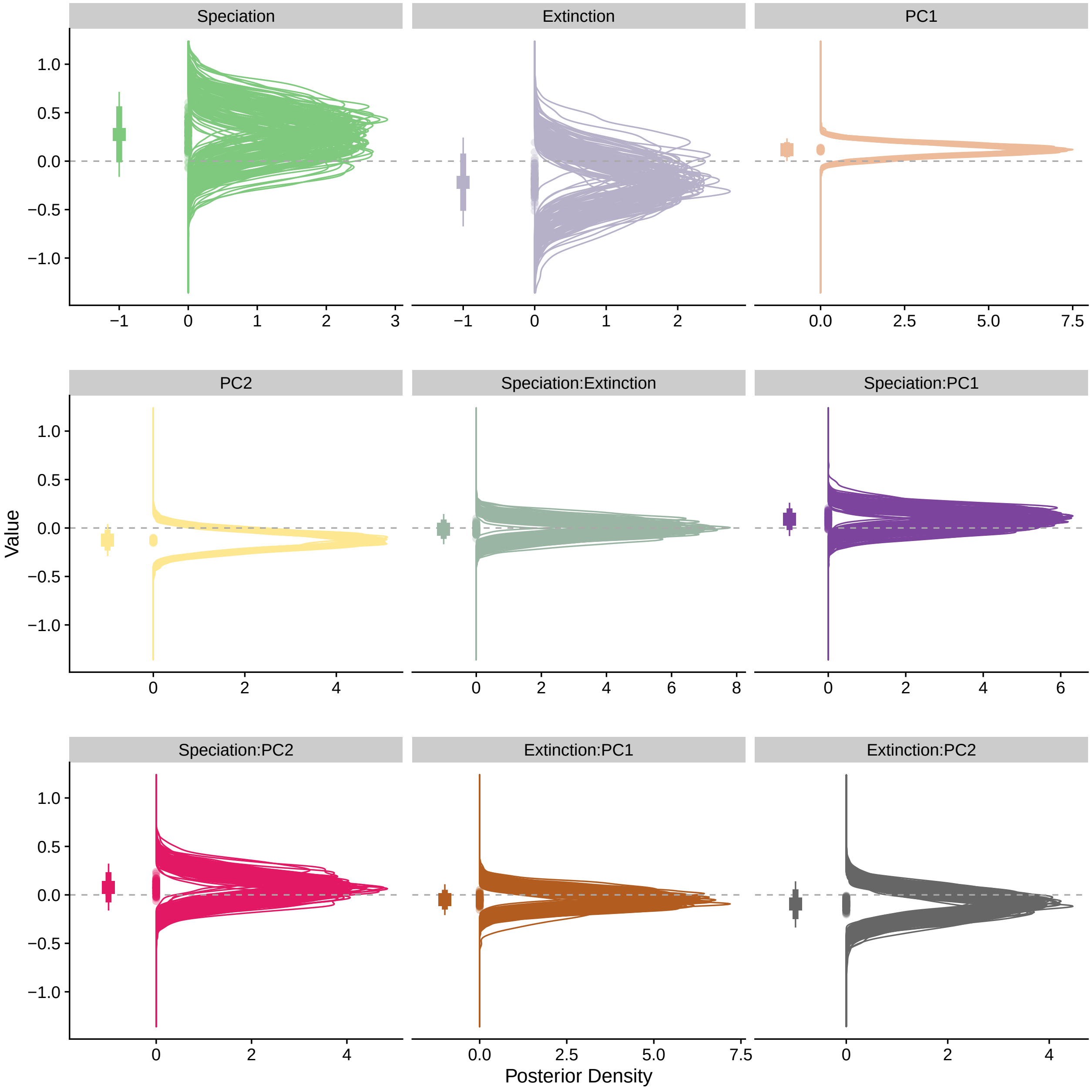


Figure S2: Individual posterior distributions for all trees using standardized speciation and extinction rates.

#### Sensitivity analysis: Individual posterior distributions per tree - Standardized Epsilon and r

The main figure shows the combined posterior distribution for all 112 trees used in the study. Below the posterior distributions for each tree are plotted individually.

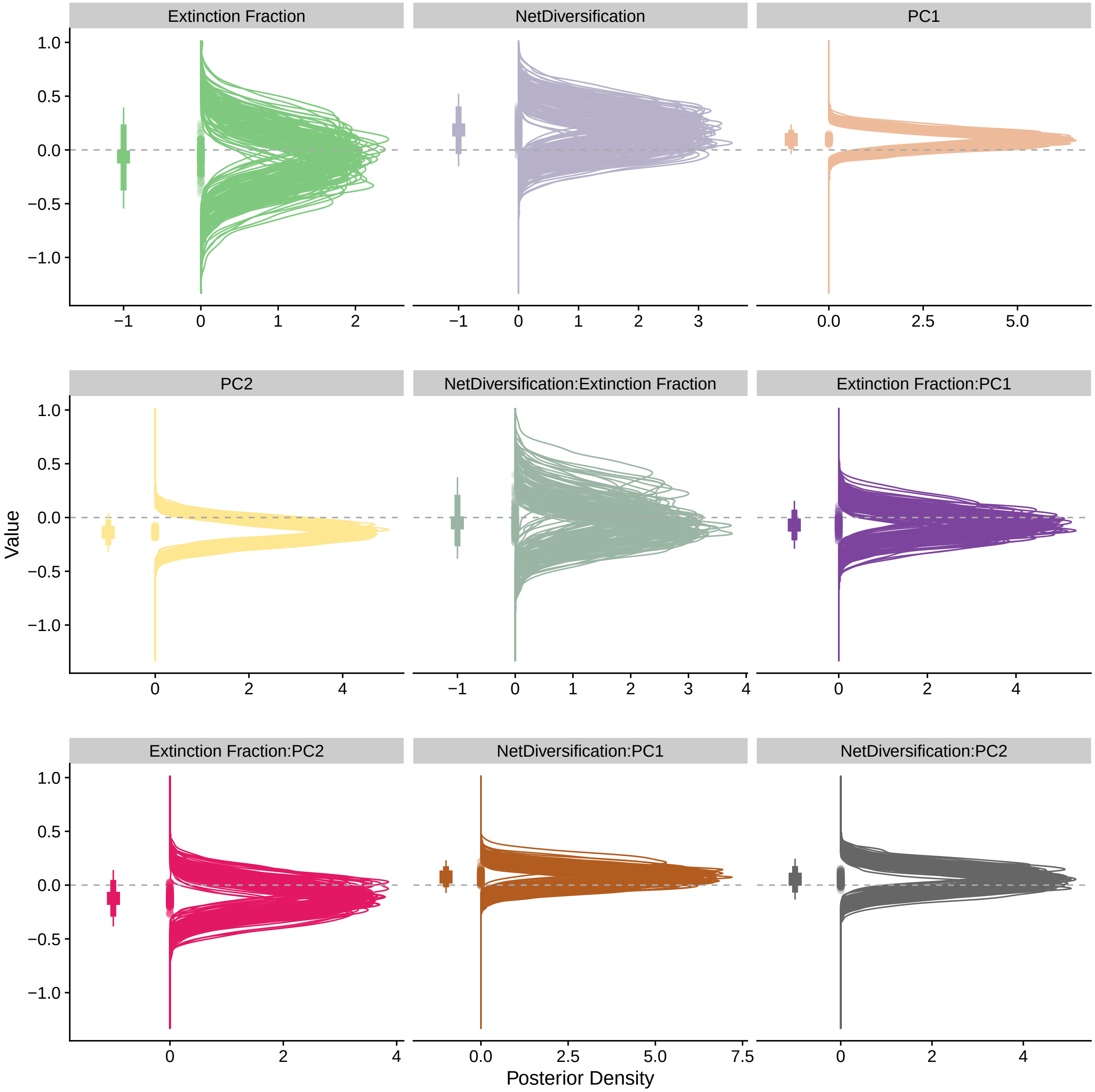


Figure S3: Individual posterior distributions for all trees used using standardized extinction fraction and net diversification rates.

#### Sensitivity analysis: Individual posterior distributions per tree - Standardized lambda and mu

The main figure shows the combined posterior distribution for all 112 trees used in the study. Below the posterior distributions for each tree are plotted individually.

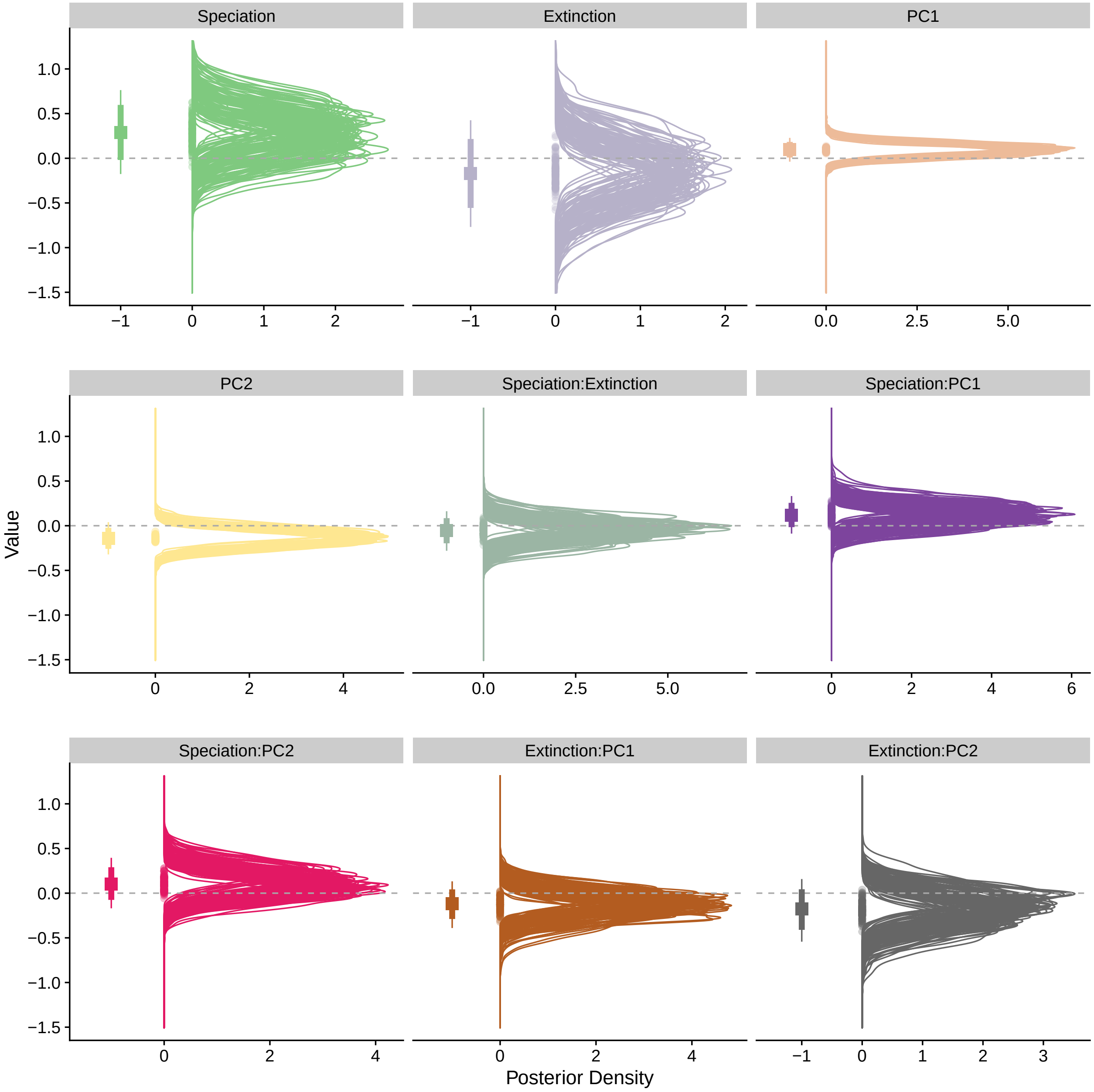


Figure S4: Individual posterior distributions for all trees using standardized speciation and extinction rates.

#### Results using raw rates - Epsilon and r

The main figure shows the combined posterior distribution for all 112 trees used in the study from the analysis using raw rates per network. Below the posterior distributions for each tree are plotted individually for the analysis using raw diversification rates.

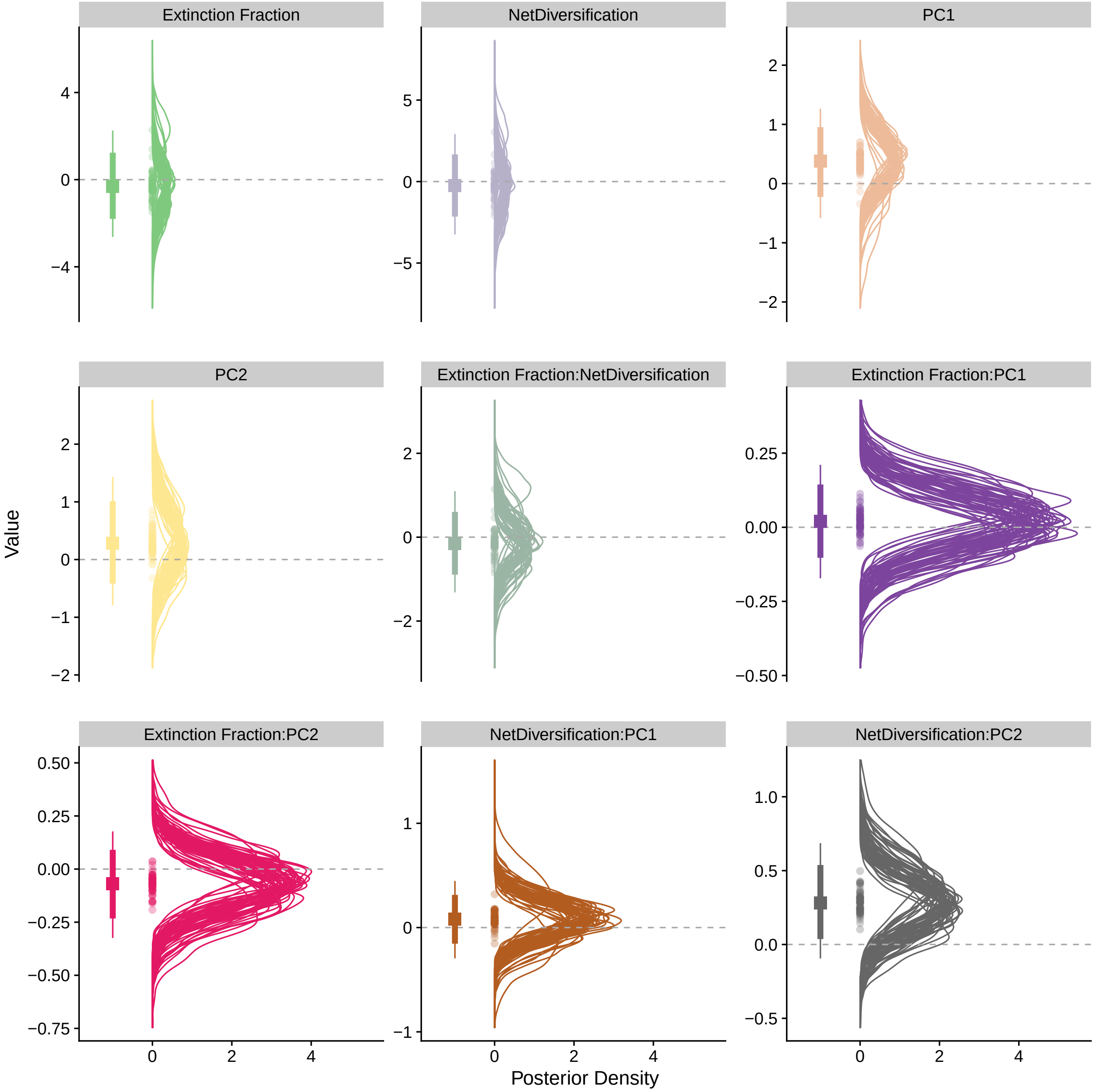


Figure S5: Posterior distributions of parameters using raw extinction fraction and net diversification rates.

#### Results using raw rates - Lambda and mu

The main figure shows the combined posterior distribution for all 112 trees used in the study from the analysis using raw rates per network. Below the posterior distributions for each tree are plotted individually for the analysis using raw diversification rates.

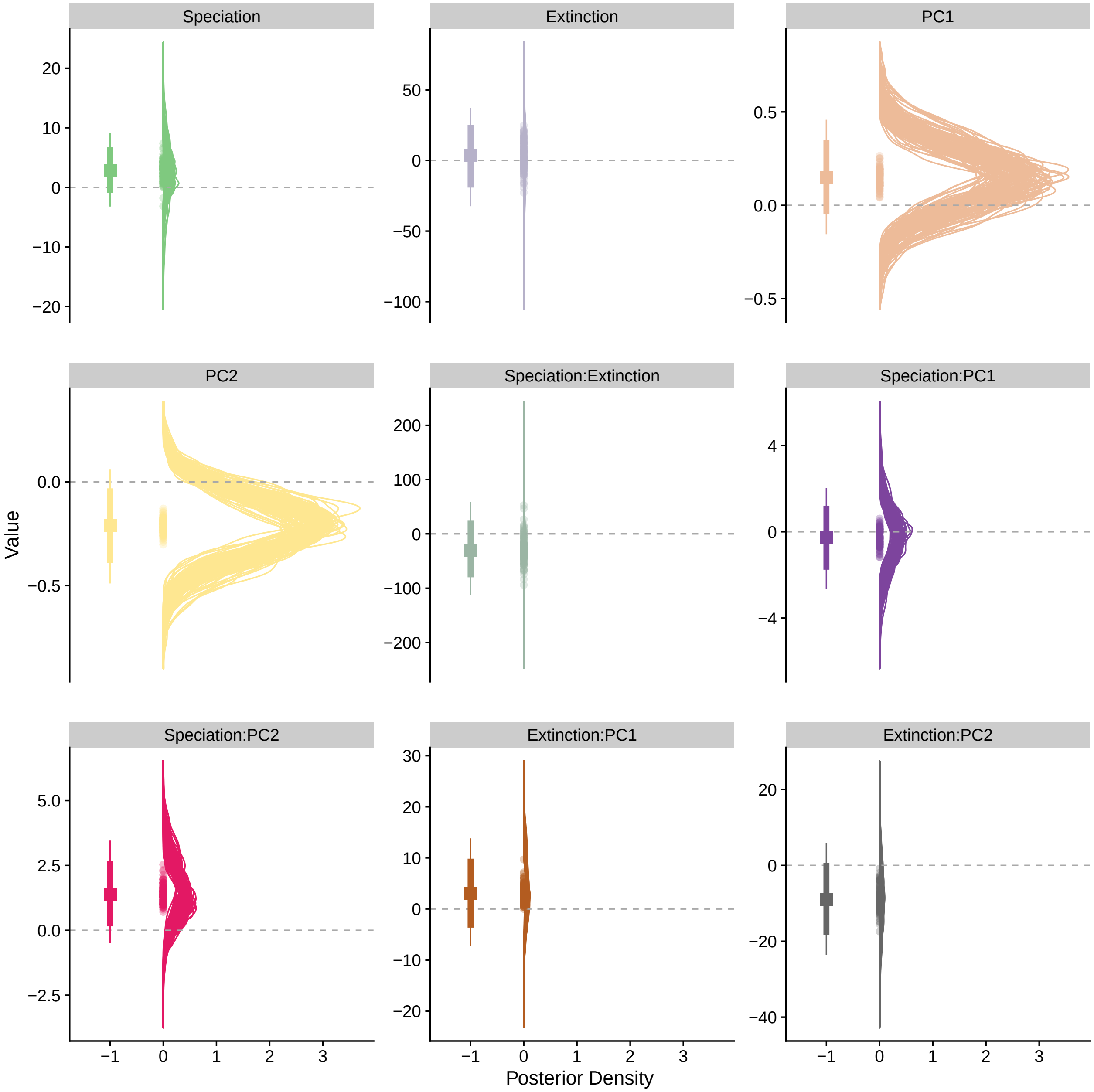


Figure S6: Posterior distributions of parameters using raw speciation and extinction rates.

#### Ecological role similarity by distance

Figure 3 in the main text shows the null distributions generated by substituting missing lineage by 0 only in the network where the lineage is not present. The plots below show the same results but with the null distribution generated substituting the mean lineage centrality in both networks by 0.

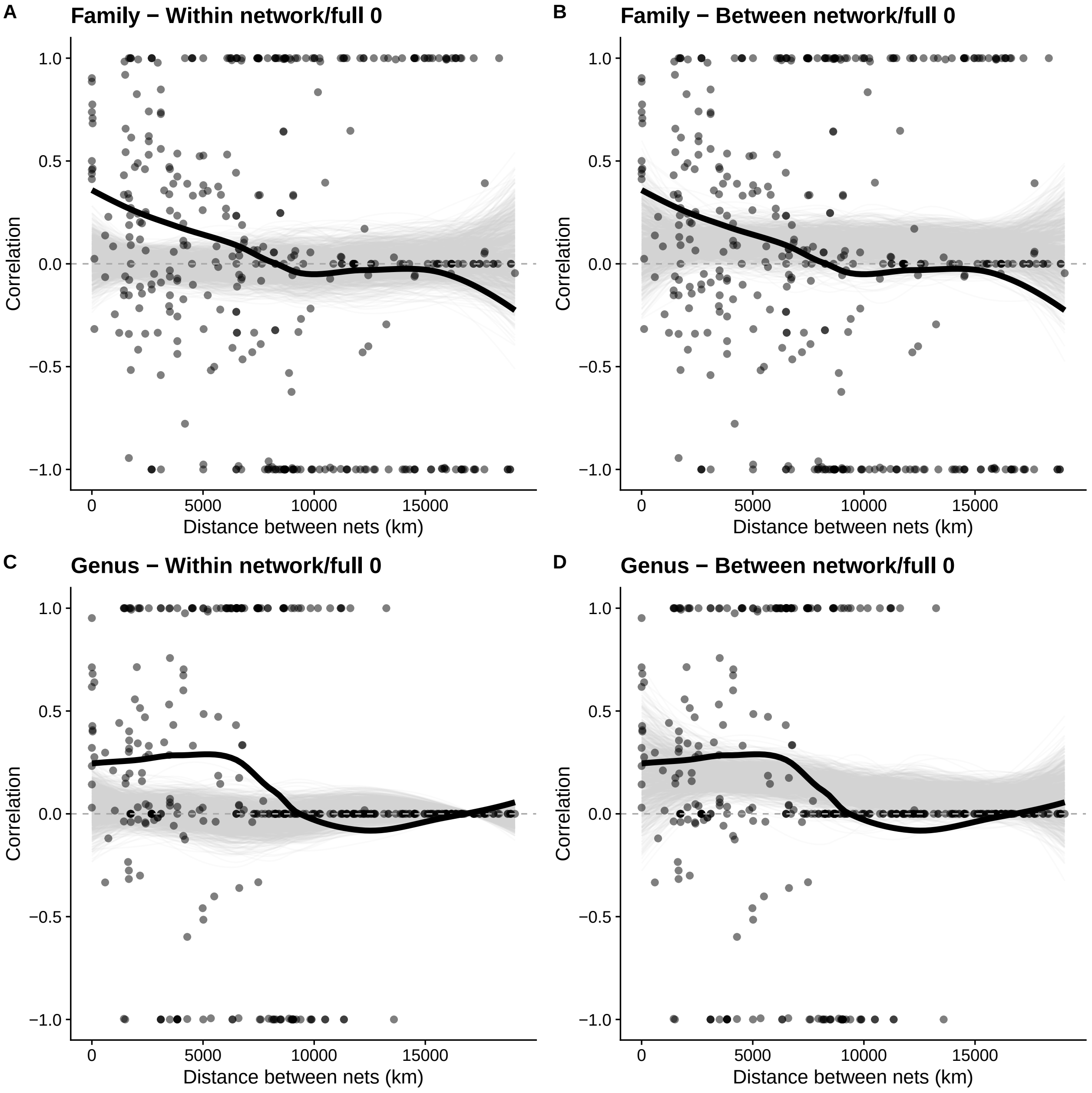


Figure S7: Association between mean PCA centrality per family (A and B) or genus (C and D) and geographical distance, using the full 0 null model where the average centrality of missing lineages is replaced by 0 in both networks.

#### Climatic PCA

Biplot of PCA of climatic factors (ann.prec = Annual Total Precipitation; prec.seas = Precipitation Seasonality; ann.mean.temp = Annual Mean Temperature; temp.seas = Temperature Seasonality).

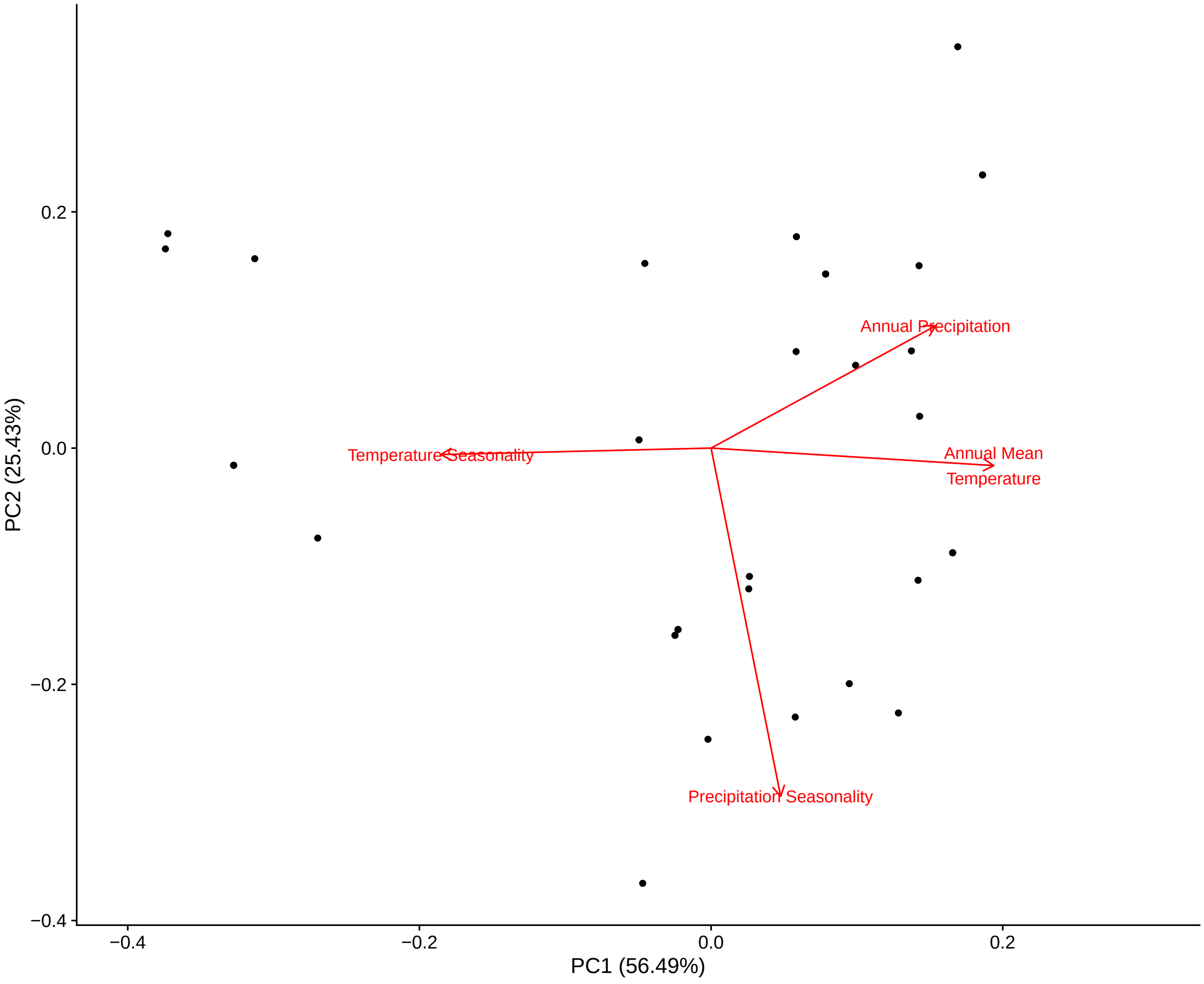


Figure S8: PCA of the climatic variables used in the study. It is worth noting that temperature variables align well with PC1 and precipitation align well with PC2 - although there is some of it on PC1 as well

#### PCA centrality

Relationship between the three centrality metrics used in the paper.

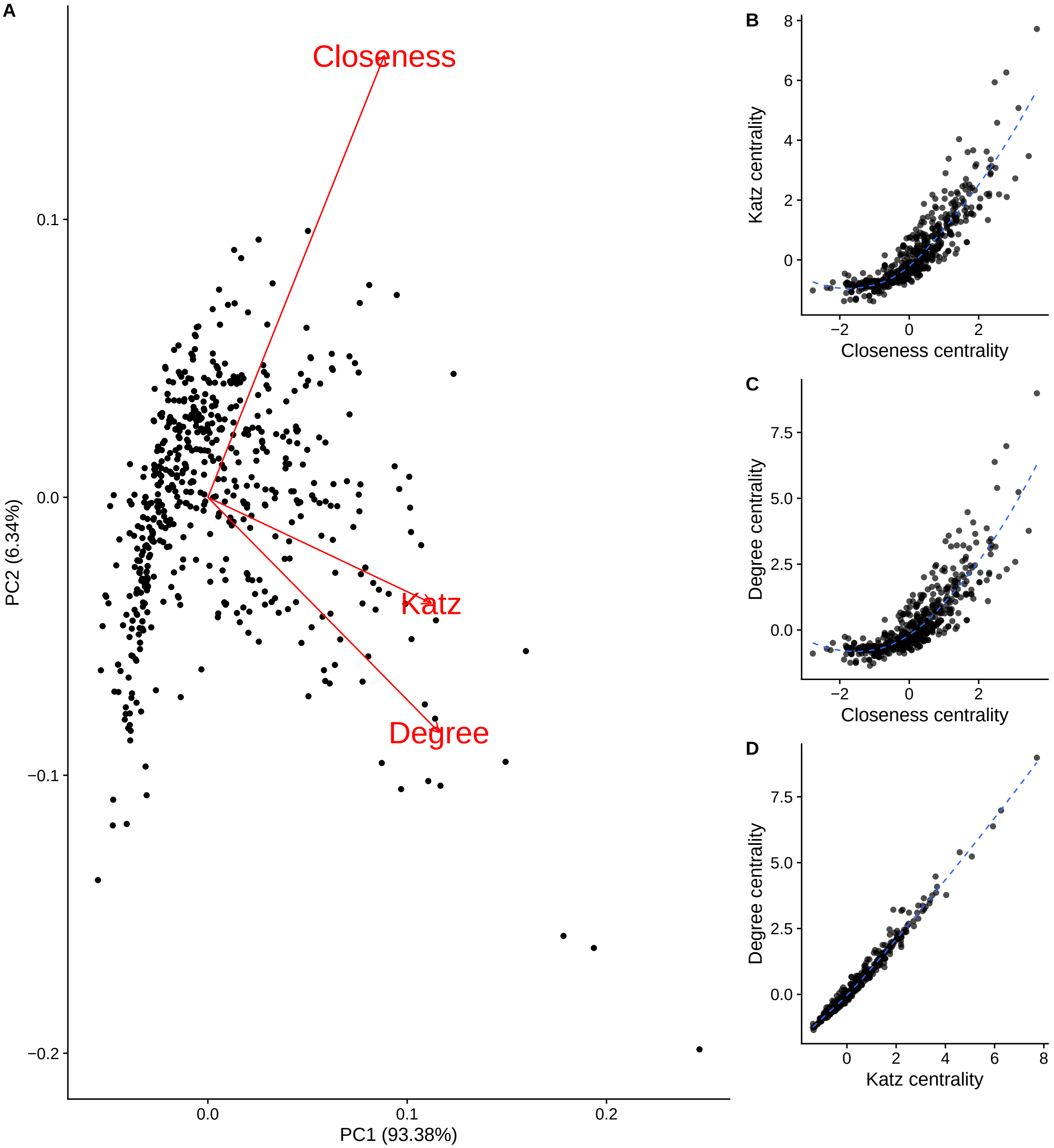


Figure S9: A: PCA of the centrality metrics (degree, closeness and katz) used in the study. The PCA was calculated using all metrics scaled per network and combined in a single dataset and PC1 was used as the general centrality since it explains 93.38% of the total variance.

### Ecospace Occupation

#### Dietary phyPCA summary

Table S2: Loadings of each dietary item in each principal component of the Phylogenetic PCA with dietary data.

| | PC1 | PC2 | PC3 | PC4 | PC5 | PC6 | PC7 | PC8 | PC9 |
|---|---|---|---|---|---|---|---|---|---|
| Invertebrates | 0.898 | -0.391 | 0.195 | -0.046 | 0.001 | 0.011 | -0.024 | 0.004 | NaN |
| Fish | -0.081 | 0.140 | -0.149 | -0.035 | 0.046 | -0.204 | 0.248 | -0.919 | NaN |
| Unknown | -0.052 | -0.034 | 0.085 | 0.044 | -0.181 | 0.541 | 0.806 | 0.104 | NaN |
| Carrion | -0.094 | -0.023 | 0.189 | 0.229 | -0.146 | -0.894 | 0.260 | 0.123 | NaN |
| Fruit | -0.881 | -0.436 | 0.175 | -0.056 | 0.000 | 0.012 | -0.026 | 0.004 | NaN |
| Nectar | -0.026 | 0.010 | -0.270 | 0.949 | 0.138 | 0.045 | -0.069 | 0.012 | NaN |
| Seeds | -0.053 | 0.946 | 0.313 | -0.054 | -0.024 | 0.016 | -0.033 | 0.006 | NaN |
| Plant Material | 0.072 | 0.048 | -0.732 | -0.221 | -0.634 | 0.024 | -0.068 | 0.013 | NaN |
| Vertebrates | 0.056 | 0.183 | -0.622 | -0.410 | 0.637 | 0.009 | -0.033 | 0.019 | NaN |

Table S3: Individual and cumulative variance explained by each of the principal components of the Phylogenetic PCA with dietary data

| | PC1 | PC2 | PC3 | PC4 | PC5 | PC6 | PC7 | PC8 | PC9 |
|---|---|---|---|---|---|---|---|---|---|
| Standard deviation | 0.071 | 0.053 | 0.030 | 0.023 | 0.020 | 0.008 | 0.006 | 0.004 | NaN |
| Proportion of Variance | 0.514 | 0.285 | 0.091 | 0.056 | 0.042 | 0.006 | 0.004 | 0.001 | 0 |
| Cumulative Proportion | 0.514 | 0.800 | 0.891 | 0.947 | 0.989 | 0.994 | 0.999 | 1.000 | 1 |

#### Summaries from linear models between disparity and climatic PCs

Summary of linear models between amplitude, average pairwise distance and mean nearest-neighbor distance and the climatic PC1.

Table S4: Summaries from linear models between disparity metrics and climatic PCs

| Disparity Metric | p-value ClimPC1 | R2 ClimPC1 | p-value ClimPC2 | R2 ClimPC2 |
|---|---|---|---|---|
| Amplitude | 0.39613 | -0.00924 | 0.22590 | 0.01878 |
| Mean NN Distance | 0.11962 | 0.05352 | 0.36031 | -0.00481 |
| Average Distance | 0.00006 | 0.43158 | 0.36933 | -0.00598 |

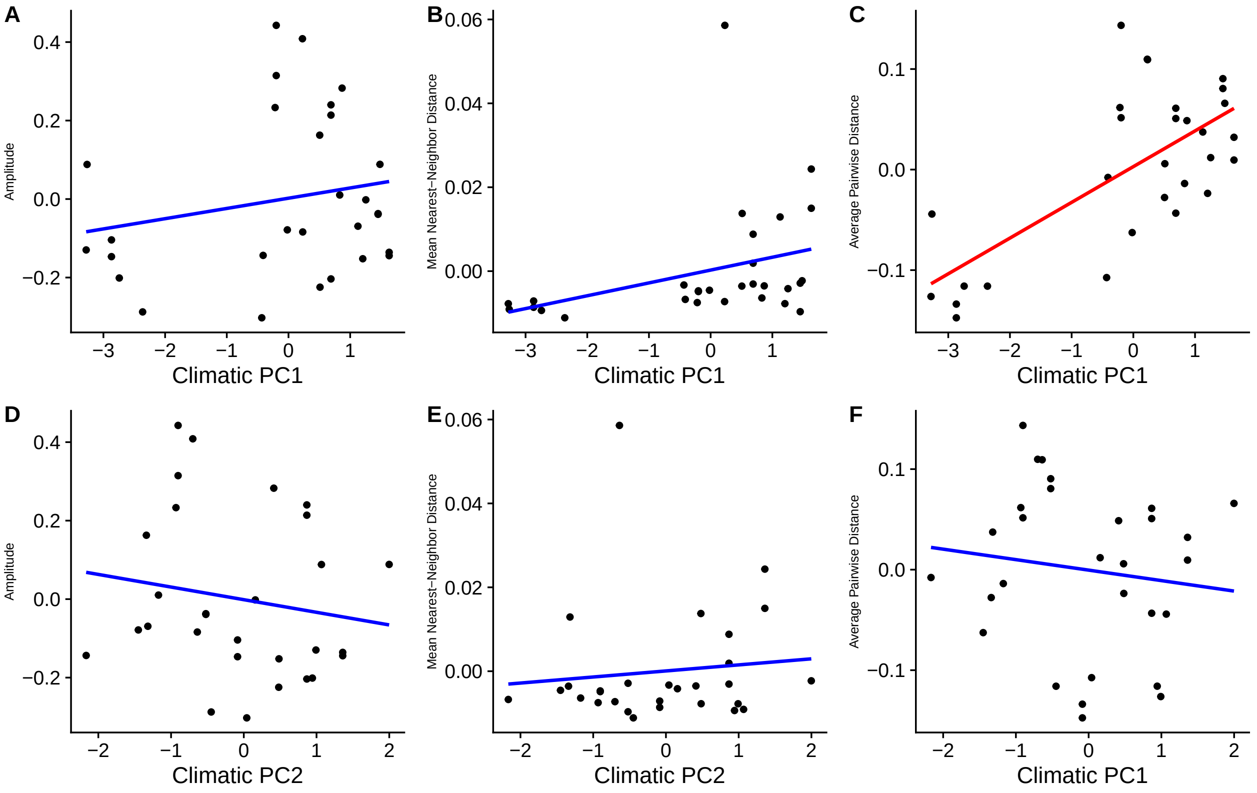


Figure S10: Different axes of specialization of bird species analyzed in the study. A - C: Relationship between Amplitude, Mean Nearest-Neighbor Distance and Average Pairwise Distance of species and Climatic PC1. D - F: Relationship between Amplitude, Mean Nearest-Neighbor Distance and Average Pairwise Distance of species and Climatic PC2. The only significant relationship is found between Average Pairwise Distance and Climatic PC1 (panel C - red line).

#### Summaries from linear models between residuals from correlation between disparity and species richness and climatic PCs

Summary of linear models between amplitude, average pairwise distance and mean nearest-neighbor distance and the climatic PC1.

Table S5: Summaries from linear models between residual disparity metrics and climatic PCs

| Disparity Metric | p-value ClimPC1 | R2 ClimPC1 | p-value ClimPC2 | R2 ClimPC2 |
|---|---|---|---|---|
| Amplitude | 0.29753 | 0.00456 | 0.40931 | -0.01074 |
| Mean NN Distance | 0.06913 | 0.08446 | 0.59103 | -0.02580 |
| Average Distance | 0.00003 | 0.46466 | 0.49624 | -0.01908 |

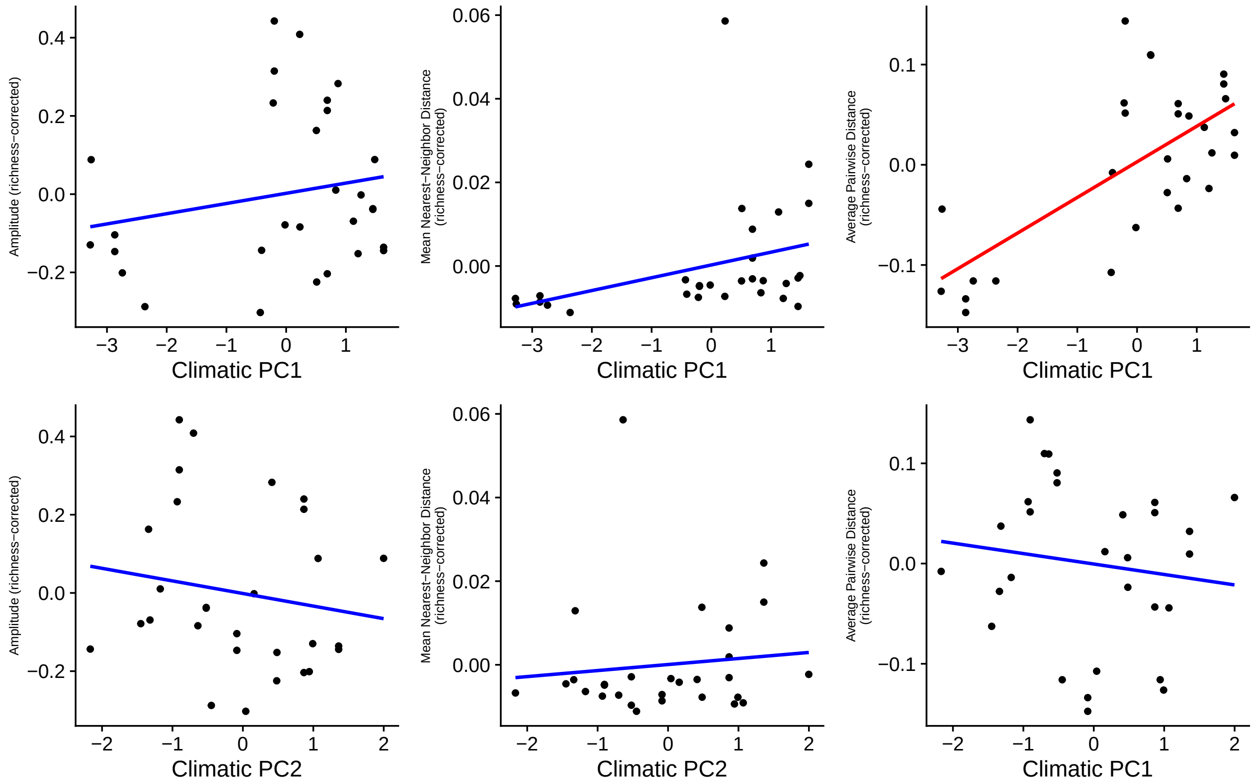


Figure S11: Different axes of specialization of bird species analyzed in the study. A - C: Relationship between Amplitude, Mean Nearest-Neighbor Distance and Average Pairwise Distance of species (accounted for species richness) and Climatic PC1. D - F: Relationship between Amplitude, Mean Nearest-Neighbor Distance and Average Pairwise Distance of species (accounted for species richness) and Climatic PC2. The only significant relationship is found between Average Pairwise Distance and Climatic PC1 (panel C - red line).

### Degree of specialization

#### Association between centrality and Number of Plant Families

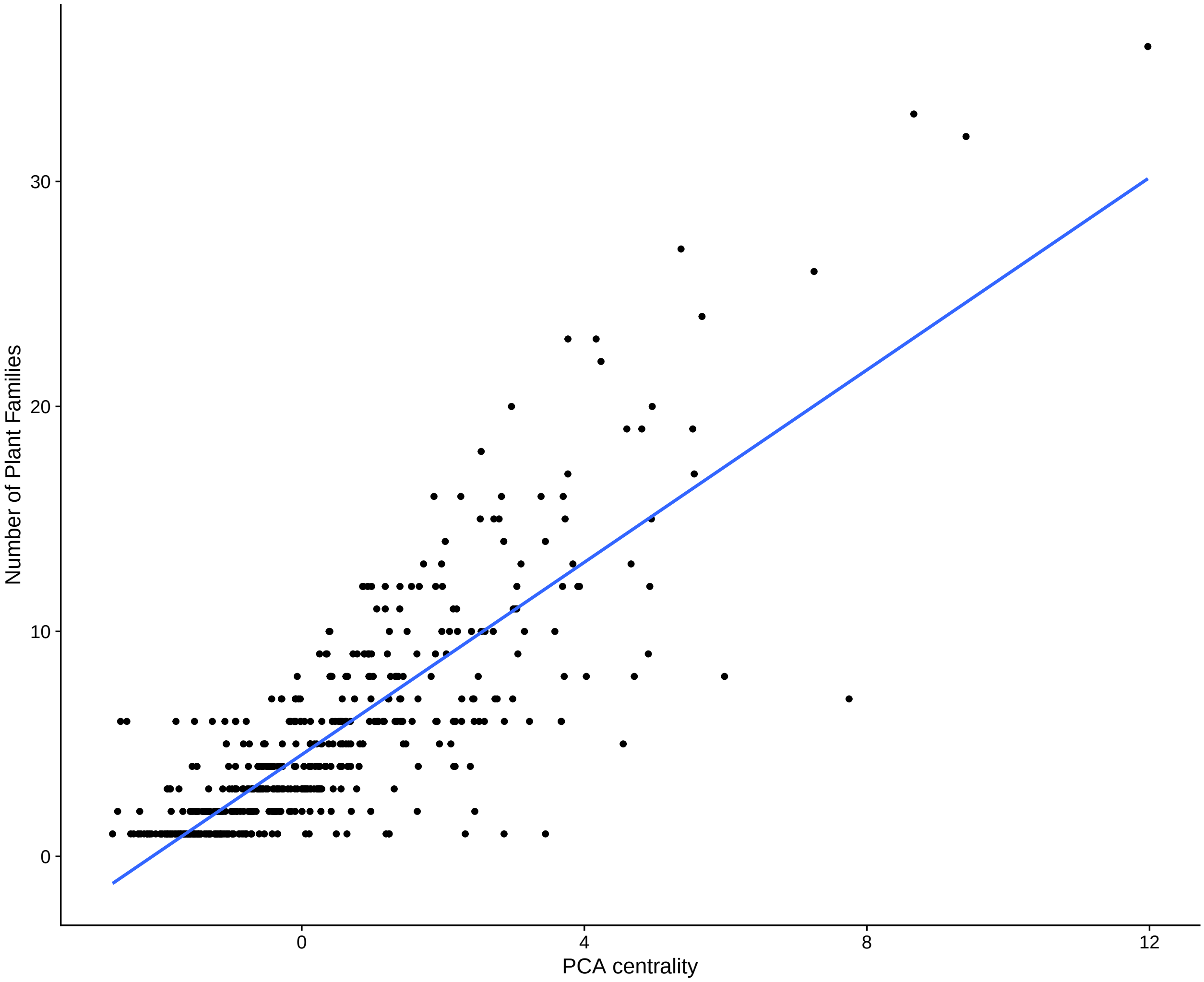


Figure S12: Relationship between PCA centrality of birds and number of plant families, showing that the more central a species is, the more generalist it is concerning the identity of their partners

Table S6: Summary of linear model testing for the relationship between PCA centrality and Number of Plant Families. $R^2$ = 0.6947

| | Estimate | Std. Error | t value | Pr(>\|t\|) |
|---|---|---|---|---|
| (Intercept) | 4.522377 | 0.1077509 | 41.97066 | 0 |
| pca | 2.137804 | 0.0578401 | 36.96057 | 0 |

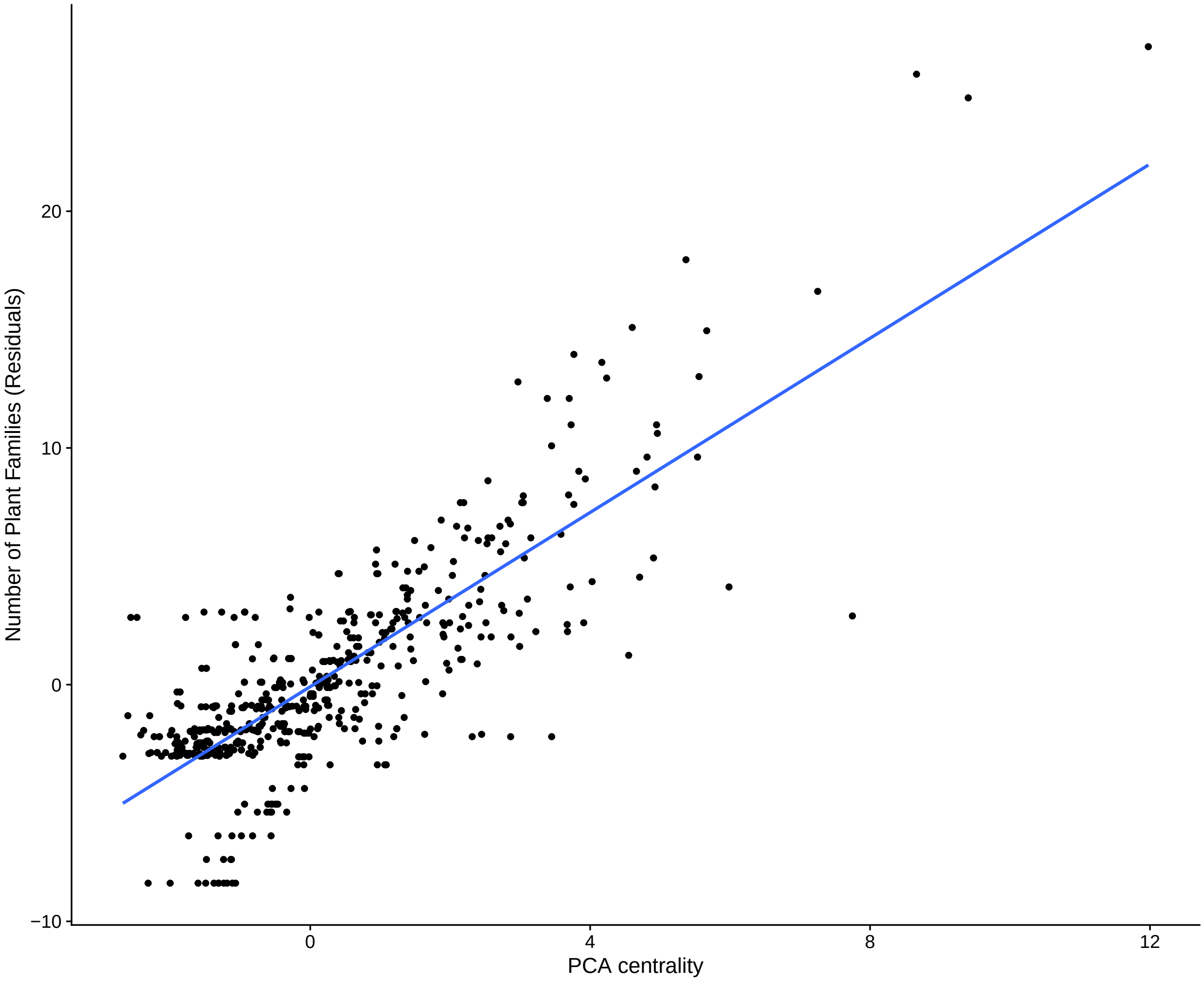


Figure S13: Relationship between PCA centrality of birds and phylogenetic diversity of plants accounting for the total number of plant species, showing that the more central a species is, the more generalist it is concerning the identity of their partners.

Table S7: Summary of linear model testing for the relationship between PCA centrality and Number of Plant Families, accounting for the total number of plant families. $R^2$ = 0.6584

| | Estimate | Std. Error | t value | Pr(>\|t\|) |
|---|---|---|---|---|
| (Intercept) | -0.086003 | 0.1007620 | -0.8535264 | 0.3937086 |
| int.tax$pca[!is.na(int.tax$n.plant.families)] | 1.840064 | 0.0540885 | 34.0195011 | 0.0000000 |

#### Association between centrality and Phylogenetic Diversity of Plants

The following tables show the summary of the association between PCA centrality and phylogenetic diversity of plants controlled either by the number of plant families or species.

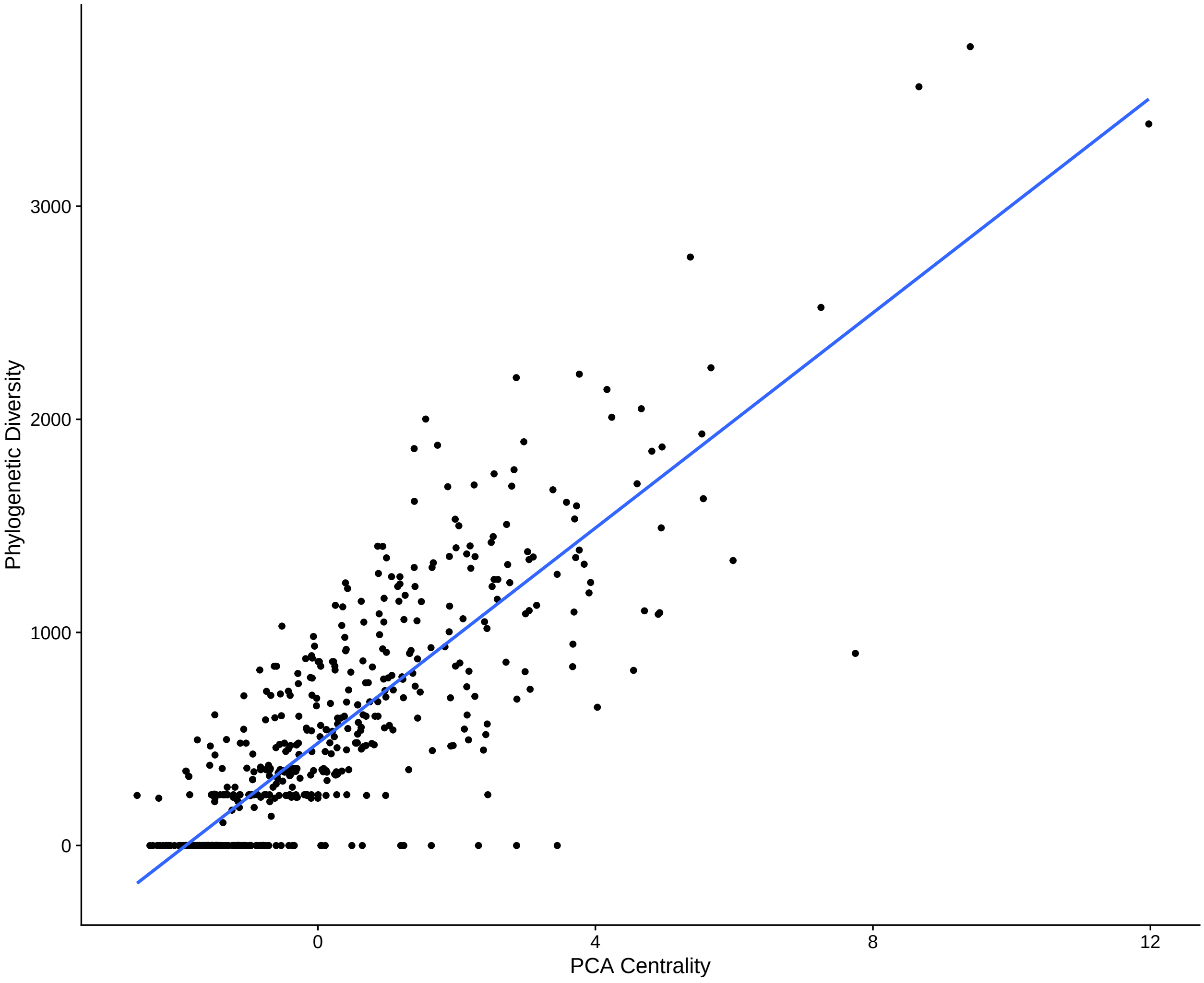


Figure S14: Relationship between PCA centrality of birds and phylogenetic diversity of plants, showing (although with a very modest fit; but see figures S12, S13, S14) that the more central a species is, the more generalist it is concerning the identity of their partners while accounting for the total number of plant families. $R^2$ = 0.0131

Table S8: Summary of linear model testing for the relationship between PCA centrality and Phylogenetic Diversity. $R^2$ = 0.7033

| | Estimate | Std. Error | t value | Pr(>\|t\|) |
|---|---|---|---|---|
| (Intercept) | 480.7704 | 12.955980 | 37.10799 | 0 |
| pca | 252.3482 | 6.880751 | 36.67452 | 0 |

Table S9: Summary of linear model testing for the relationship between PCA centrality and Phylogenetic Diversity, accounting for the total number of families

| | Estimate | Std. Error | t value | Pr(>\|t\|) |
|---|---|---|---|---|
| (Intercept) | -1.10847 | 7.503556 | -0.147726 | 0.8826116 |
| int.tax$pca[!is.na(int.tax$phylodiv)] | 11.63989 | 3.985040 | 2.920896 | 0.0036294 |

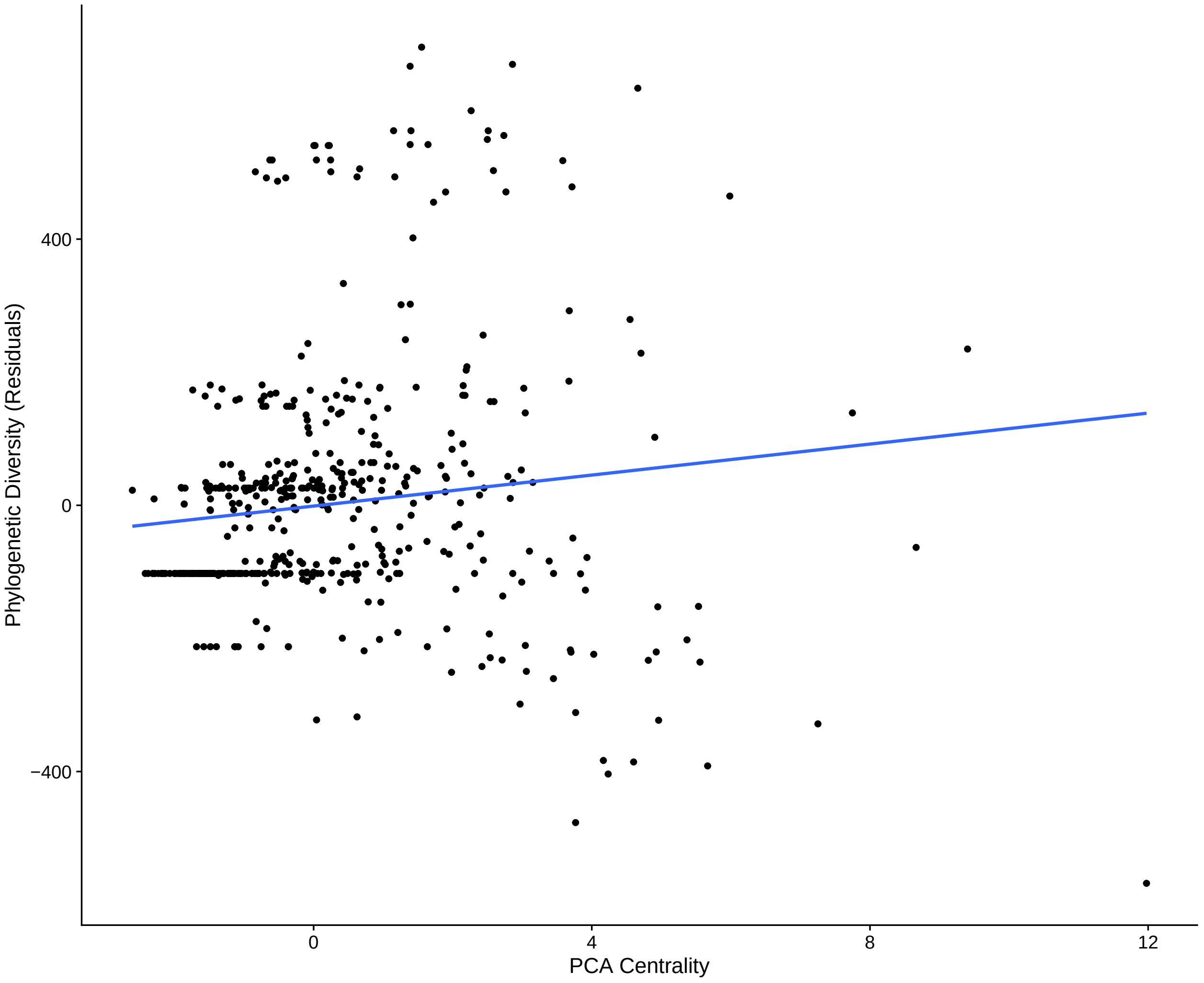


Figure S15: Relationship between PCA centrality of birds and phylogenetic diversity of plants accounting for the total number of plant families, showing that the more central a species is, the more generalist it is concerning the identity of their partners. $R^2$ = 0.0131
